# Proteome-wide *in silico* screening for human protein-protein interactions

**DOI:** 10.1101/2025.11.10.687652

**Authors:** Ernst W. Schmid, Helen Zhu, Eunjin Ryu, Yang Lim, Agata Smogorzewska, Alan Brown, Johannes C. Walter

## Abstract

Protein-protein interactions (PPIs) drive virtually all biological processes, yet most PPIs have not been identified and even more remain structurally unresolved. We developed a two-step computational screen for human PPIs. First, a classifier called KIRC (Knowledge-Informed Rapid Classifier), trained on biological features, was used to rank all 200 million possible protein pairs in the human proteome by their interaction likelihood. Second, the ∼1.6 million top-ranked KIRC pairs were subjected to structure prediction by AlphaFold-Multimer and ranked using SPOC (Structure Prediction and Omics Classifier), which identifies functional predictions based on biological and structural features. This pipeline revealed 16,000 high-confidence PPIs (∼90% precision), of which more than 5,000 were not previously recognized and more than 12,000 have not been structurally resolved. We use this “predictome” to formulate new hypotheses in different areas of biology, reinterpret low-resolution cryo-EM maps, and identify and validate novel PPIs that may support replication-coupled chromatin assembly. The predicted PPIs, viewable at predictomes.org, are expected to accelerate characterization of the molecular interactions that underlie vertebrate cell physiology.

## Introduction

The human genome encodes ∼20,000 proteins, most of which engage in protein-protein interactions (PPIs). Stable PPIs generate complex cellular structures and machines, whereas transient interactions underlie dynamic processes such as DNA replication and cell signaling^1^. How many of the 200 million (M) possible binary protein combinations comprise functional PPIs is not known, with estimates ranging from 70,000 to 650,000^2^. Only ∼9,000 of these PPIs have been resolved structurally, leaving a large gap in our understanding of the interactions that execute cellular functions.

In 2022, DeepMind developed AlphaFold-Multimer (AF-M), a deep learning system trained to predict the structure of protein complexes^3^. Importantly, AF-M reports its confidence in each prediction, a feature we and others used to identify new PPIs. In this “*in silico*” screening approach, a bait protein is “folded” with different prey proteins, and the resulting binary structure predictions are ranked by confidence^4–7^. In several cases, the top ranked PPIs were subsequently shown to represent physiologically relevant interactions based on structure-guided mutagenesis and functional analysis^6,8–12^ . For example, *in silico* screening identified functional binding partners of proteins involved in DNA replication^6^, transcription-coupled nucleotide excision repair^11^, and fertilization^12^. In every case, the newly identified interaction led to novel mechanistic insights. Thus, *in silico* screening represents a powerful approach to discover, structurally resolve, and functionally dissect PPIs, thereby dramatically accelerating our understanding of molecular mechanisms.

An exciting prospect is to use *in silico* screening to identify all human binary PPIs (the human “predictome”). This goal faces two challenges. First, AF-M confidence metrics are imperfect and lead to many false positives^13^. To identify physiologically relevant AF-M predictions, we recently trained a “classifier” called SPOC (Structure Prediction and Omics Classifier) that assesses whether a binary prediction is both structurally plausible *and* consistent with experimental omics data such as protein co-localization, co-precipitation, and genetic co-dependency^14^. We showed that SPOC strongly enriches for functional protein pairs and thus enables proteome-wide *in silico* screening^14^. The second barrier to generating a comprehensive predictome is the availability of computational resources. Folding all 200 M human protein pairs with AF-M would require ∼50 M graphics processing unit (GPU) hours. Therefore, before attempting proteome-wide structure prediction, a triage step is currently needed to nominate likely interactors for analysis by AF-M.

To identify potential PPIs, we trained a new classifier called KIRC (Knowledge Integrated Rapid Classifier) that scores the likelihood that two proteins interact based on features derived from biological omics data. Because KIRC is computationally inexpensive, we could calculate KIRC scores for all 200 M human protein pairs in 24 hours. The 1.6 M pairs with the highest KIRC scores then underwent structure prediction using AF-M and scoring by SPOC. This screen yielded more than 16,000 high-confidence heterodimeric PPIs at ∼90% precision. 3,495 of these PPIs share sequence similarity to previously determined interactions in the protein databank (PDB), leaving a total of 13,056 pairs with no existing homologous experimental structures. Of these, 5,084 have no strong prior evidence of interaction, and are therefore considered novel. Data for all 1.6 million pairs can be browsed interactively at predictomes.org. The predictome facilitates the interpretation of low-resolution cryo-EM maps, and enables the formulation and testing of novel hypotheses, including that the DNA replication factor DONSON participates in chromatin assembly. The identification of thousands of new, structurally resolved PPIs is expected to accelerate mechanistic understanding across cell physiology.

## Results

### A new classifier for rapid triage of potential PPIs

Analyzing all 200 M possible human protein pairs using AF-M is not currently feasible given that each prediction requires several minutes of compute on a GPU. We therefore asked whether a less complex classifier trained on biological features alone could be used as a rapid triage step to nominate pairs for full analysis by AF-M and SPOC. To address this, we complemented the biological features originally used to train SPOC with additional features such as whether two proteins correlate in co-fractionation mass spectrometry data (See Methods). We also included a recently reported deep learning-based interaction probability (RF2-PPI) score^15^.

Using this expanded set of biological features (**Table S1**), we trained a new random forest classifier to nominate interacting protein pairs. For training, we curated two positive data sets that contain interacting pairs and four negative sets containing non-interacting pairs (**Table S2** and Methods)^14^. To optimize performance, we systematically varied the balance of these six datasets during training (hyperparameter tuning), generating thousands of distinct classifiers (**Table S3**). We tested their performance by assessing how they ranked a bait protein’s interacting partner (not in the training set) in a list of all ∼20,000 human proteins, the vast majority of which are not interactors (**Figure 1A**). In 39 such “ranking experiments” (**Table S4**), the highest performing classifier achieved a median rank of 23 and placed 31 out of 39 true pairs within the top 100 hits (**Figure 1B**). For comparison, STRINGDB^16^ assigned a median rank of 55, and only 26 true pairs ranked within the top 100 (**Figure 1B**).

**Figure 1:**
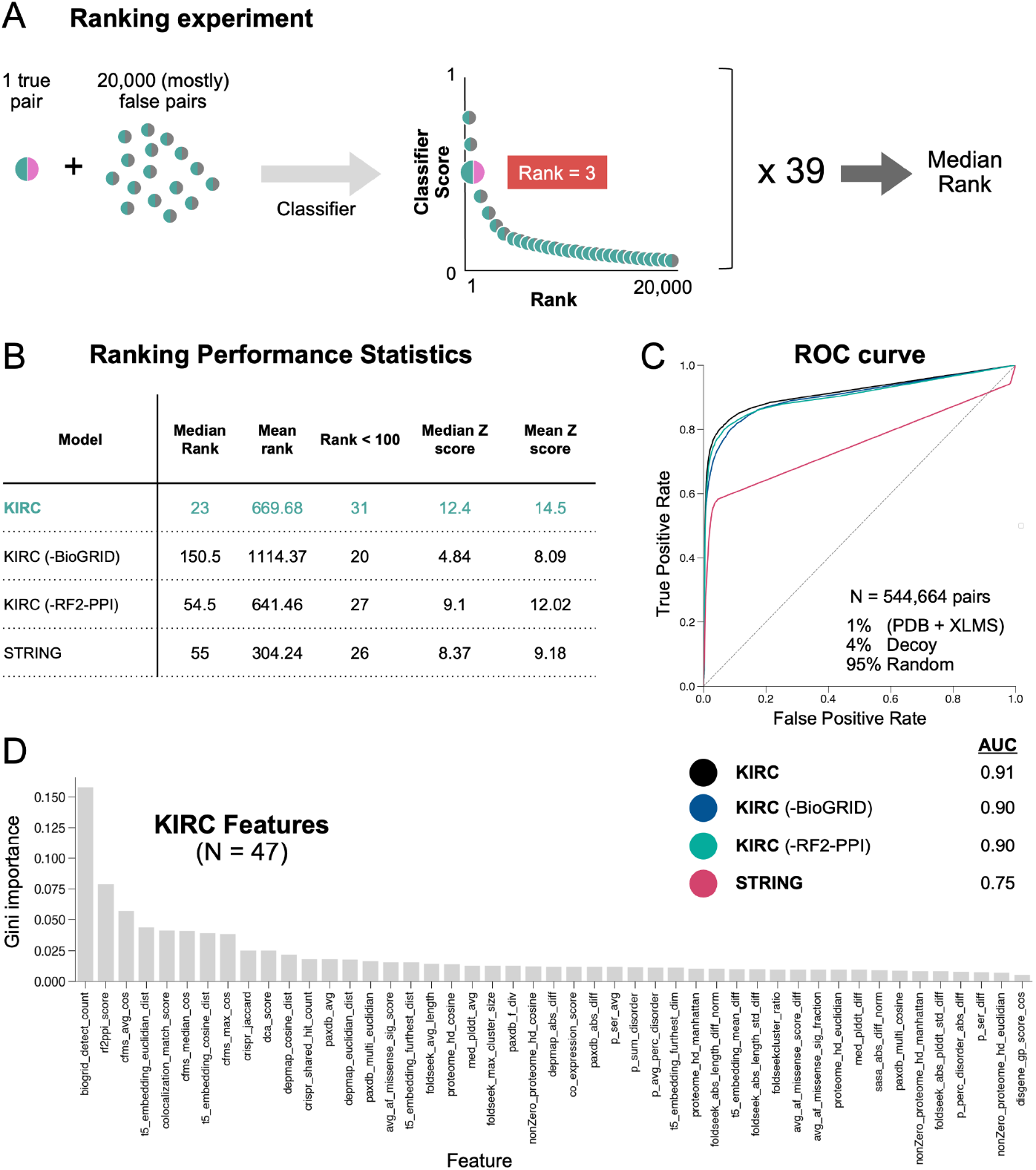
Developing a classifier for rapid nomination of likely protein interactors. **A**. Schematic illustrating how proteome wide ranking experiments were used to assess classifier performance. The procedure was repeated for 39 benchmark pairs, and the median rank was calculated. **B**. A table comparing the performance of various metrics and classifiers on the 39 ranking experiments. **C**. A Receiver Operating Characteristic (ROC) plot that compares the true positive and false positive rates associated with different metrics and classifiers at different cutoffs. Area under the curve (AUC) values are displayed for each model. An AUC of 1.0 indicates perfect discrimination, while an AUC of 0.5 is no better than random chance. **D**. Histogram of all 47 KIRC features ranked by their Gini importance values from high to low.

We additionally evaluated the classifier on ∼550,000 curated positive and negative pairs (1:99 P:N ratio) that were excluded from training, which yielded an area under the curve (AUC) of 0.91 (**Figure 1C**). The number of datapoints supporting interaction in BioGRID (“evidence count”), was the most important feature contributing to performance, followed by RF2-PPI (**Figure 1D**). Dropout of individual features showed that BioGRID was an important driver of performance, especially in ranking experiments, whereas the RF2-PPI score contributed less (**Figure 1B and 1C**). Ultimately, a classifier that includes all 47 features listed in **Table S1** was used to nominate protein pairs for full analysis by AF-M and SPOC (next section). We call this classifier KIRC (<u>K</u>nowledge-Integrated <u>R</u>apid <u>C</u>lassifier).

### Modeling 1.6 million KIRC-nominated pairs with AF-M

We next used KIRC to score the interaction likelihood of all ∼200 M human protein pairs, which took only ∼24 hours. The top ∼1.6 M heterodimeric pairs (KIRC ≥ 0.088) were then modeled by AF-M. We excluded 64,533 pairs whose combined length exceeded 3,600 residues or whose sequences were redundant with other nominated pairs. To reduce computation, each pair was folded in three out of five AF-M models using the ColabFold implementation of AF-M^17^. Pairs were folded on an NVIDIA DGX server containing 256 A100 GPUs, and MMSeqs2 was run on the central processing units (CPUs) associated with each GPU node so that multiple sequence alignment (MSA) generation was not rate-limiting (**Figure S1A-B**). Using this configuration, the 1.6 M pairs were modeled in three months (∼500,000 GPU hours). Among these, we retained only the 525,907 pairwise predictions (32.8%) in which the two chains met minimum contact criteria (“C+”: at least 5 interchain residue pairs were predicted with confidence by AF-M; see Methods). We then calculated SPOC scores for all C+ pairs and obtained the distribution shown in **Figure 2A**. To estimate the False Discovery Rate (FDR), we determined how many negative pairs were incorrectly labeled as true at each SPOC threshold. We performed this analysis on a curated dataset containing a 1:1000 P:N ratio to approximate the human interactome (∼200,000 real PPIs among 200 M possible pairs), with negative pairs being selected at random. As shown in **Figure 2B**, SPOC > 0.86 was associated with a 10% FDR (90% precision). Out of the 525,907 C+ pairs, 16,469 had a SPOC score greater than 0.86, constituting a high-confidence “16k set”. Lowering the SPOC threshold to 0.5 yielded 113,625 pairs at a 50% FDR. Among the 39 pairs used for the rank tests, 30 earned a SPOC score greater than 0.5, and 14 scored above 0.86 (**Table S4**). Our procedure likely over-estimated the FDR because some random pairs incorrectly labeled as false (false negatives) would earn high SPOC scores, inflating the FDR. Importantly, 0.86 should not be interpreted as a strict cutoff as predictions with SPOC scores below 0.86 are often correct (as demonstrated below).

**Figure 2:**
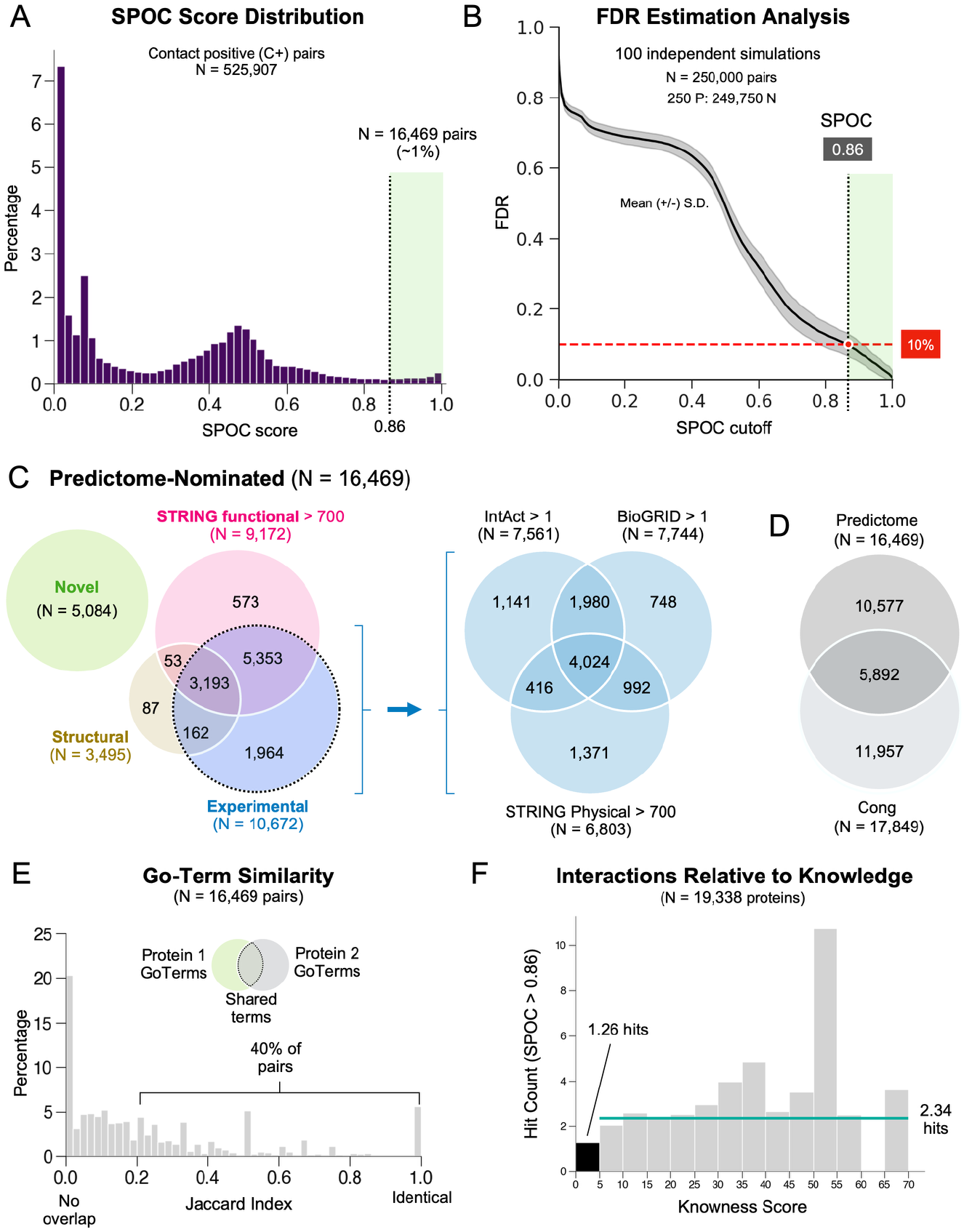
Evaluating the results of the proteome wide AF-M screen. Histogram of SPOC scores for all 525,907 contact positive “C+” pairs that achieved five or more confident contacts in at least one of the three AlphaFold-Multimer predictions. A subset of 16,469 pairs surpasses the SPOC=0.86 cutoff. Graph illustrating the False Discovery Rate (FDR) versus SPOC cutoffs in the screen as determined via simulation (see Methods). The black line displayed is the mean from 100 independent simulations. A SPOC score of 0.86 was associated with a FDR of 10% and is denoted by the dashed vertical line. **C**. Venn diagrams showing how many pairs from the high-confidence 16k set were novel versus being previously identified in STRING (functional), the PDB, or experimental databases (BioGRID, IntAct, and STRING physical). **D**. Venn diagram comparing the final high-confidence pairs identified via our screen vs those identified by a separate proteome-wide screen^15^. **E**. Histogram of GO-term similarity (Jaccard index) among proteins within pairs in the 16k set. **F**. Histogram of mean number of SPOC hits associated with proteins binned by their knowness score. The teal line is the average hit rate per protein among proteins with knowness scores >5.

### Comparison with existing databases

To address how many pairs from the 16k set were already known, we first asked how many resembled structures in the PDB. Using MMSeqs2 (ref.^18^), we mapped all unique protein pairs in the PDB to their most homologous human counterparts (minimum 30% sequence identity). This revealed 24,755 unique human protein pairs that have an identical (∼9,000) or a homologous (∼15,000) pair in the PDB (See Methods and **Figure S2A-B**), 3,495 of which were represented in the 16k set (**Figure 2C**). Thus, 21% of the confidently predicted pairs (3,495/16,469) displayed homology to known protein structures. We further addressed which predicted pairs were associated with experimental evidence in PPI databases. Based on a criterion of two or more evidence counts in IntAct or BioGRID, or a STRING physical score greater than 700, 10,672 pairs in the 16k set (64%) were supported by previous experimental data (**Figure 2C**). Our set also included 573 pairs with high *functional* STRING scores (>700) but no other evidence. In sum, 11,385 PPIs in the 16k set were supported by prior experimental, structural, or functional evidence. This leaves 5,084 high-confidence pairs with no strong prior evidence of interaction, which we designate as “novel” (**Figure 2C** and **Table S5**).

Finally, we quantified how many pairs in the 16k set overlapped with the recent proteome-wide *in silico* screen from Cong and colleagues^15^. Their pipeline yielded 17,849 pairs at 90% precision, 3,631 of which were deemed novel based on criteria analogous to those we applied above. Roughly one third of the high confidence pairs obtained (5,892 pairs) were shared between the screens (**Figure 2D**), indicating substantial agreement. However, each method also identified a unique subset of interactions, and ours produced a 40% higher yield of novel pairs (5,084 vs. 3,631).

### Well-studied proteins generally have more predicted interactors

We next examined whether our dataset was biased towards proteins involved in specific biological pathways. The 16k set includes 8,199 unique proteins, representing 41% of the canonical human proteome. Gene ontology (GO) term enrichment analysis^19^ of these proteins revealed moderate enrichment (up to 2.5x) for many pathways, with a smaller number of pathways being under-represented (**Figure S2C**). We also analyzed the relationship between GO-terms associated with proteins *within* pairs. In the 16k set, 40% of paired proteins shared the same or similar GO terms (**Figure 2E**; Jaccard ≥ 0.2), with representation across a wide range of pathways. However, for more than 20% of the confident pairs, no GO terms were shared, suggesting that in these cases, our screen identified interactions between pathways normally considered to be unrelated (**Figure 2E**).

We also asked whether the likelihood of finding confident interactors correlated with the degree to which a protein has been characterized. We first consulted the “unknome” database, which quantifies knowledge about every human protein^20^. Although “knowness” scores range from 0 to 170, many well-studied proteins such as CDC45 and NUP93 have scores of only 10. We therefore determined the average number of high-confidence hits (SPOC > 0.86) for proteins with a knowness score below 5 (poorly characterized) versus those with scores above 5 (well characterized). As shown in **Figure 2F**, poorly characterized proteins had an average of 1.26 high-confidence interactors, whereas well-characterized proteins averaged 2.34. Similarly, proteins associated with 10 or fewer publications in PubMed (N = 3,053 proteins)^21^ retrieved an average of 0.5 high-confidence SPOC hits compared to ∼2 high-confidence hits for proteins with more than 10 publications (N = 14,874 proteins; **Figure S2D**). Finally, when we analyzed SPOC hit counts as a function of UniProt annotation score (1–5 scale), well-characterized proteins were much more likely to have interactions in the 16k set (**Figure S2E**). Collectively, these results show that the PPIs in our database have no strong pathway bias but are weighted towards well-studied proteins. Nevertheless, there are 1,554 high-confidence partners predicted for the 420 most understudied proteins (publication count <10) in our high-confidence dataset. Moreover, as illustrated below, the database contains many informative new predictions for well-studied proteins.

### A web portal for exploring predicted PPIs

Our predictions are publicly available on predictomes.org^14^, a web platform that facilitates assessment of structure prediction data. Users can search the database for any human protein and retrieve all its candidate interactors, ranked by SPOC score, from the list of 525,907 C+ pairs (**Figure 3A**). In addition to the SPOC score and its associated FDR (from **Figure 2B**), the table also reports AF-M confidence metrics, data **A**. from BioGRID and STRING, and any homologous structures in the PDB, reducing the need for manual cross-referencing. Clicking on a hit reveals an information page with an interactive structure viewer that allows superposition of any homologous pairs from the PDB onto the structure prediction (**Figure 3B**). The information page also displays PAE heatmaps (**Figure S3A**), residue-level pLDDT plots (**Figure S3B**), and residue-level conservation plots (**Figure S3C**). Together, the data on predictomes.org facilitates triage of structure predictions and molecular hypothesis building.

**Figure 3:**
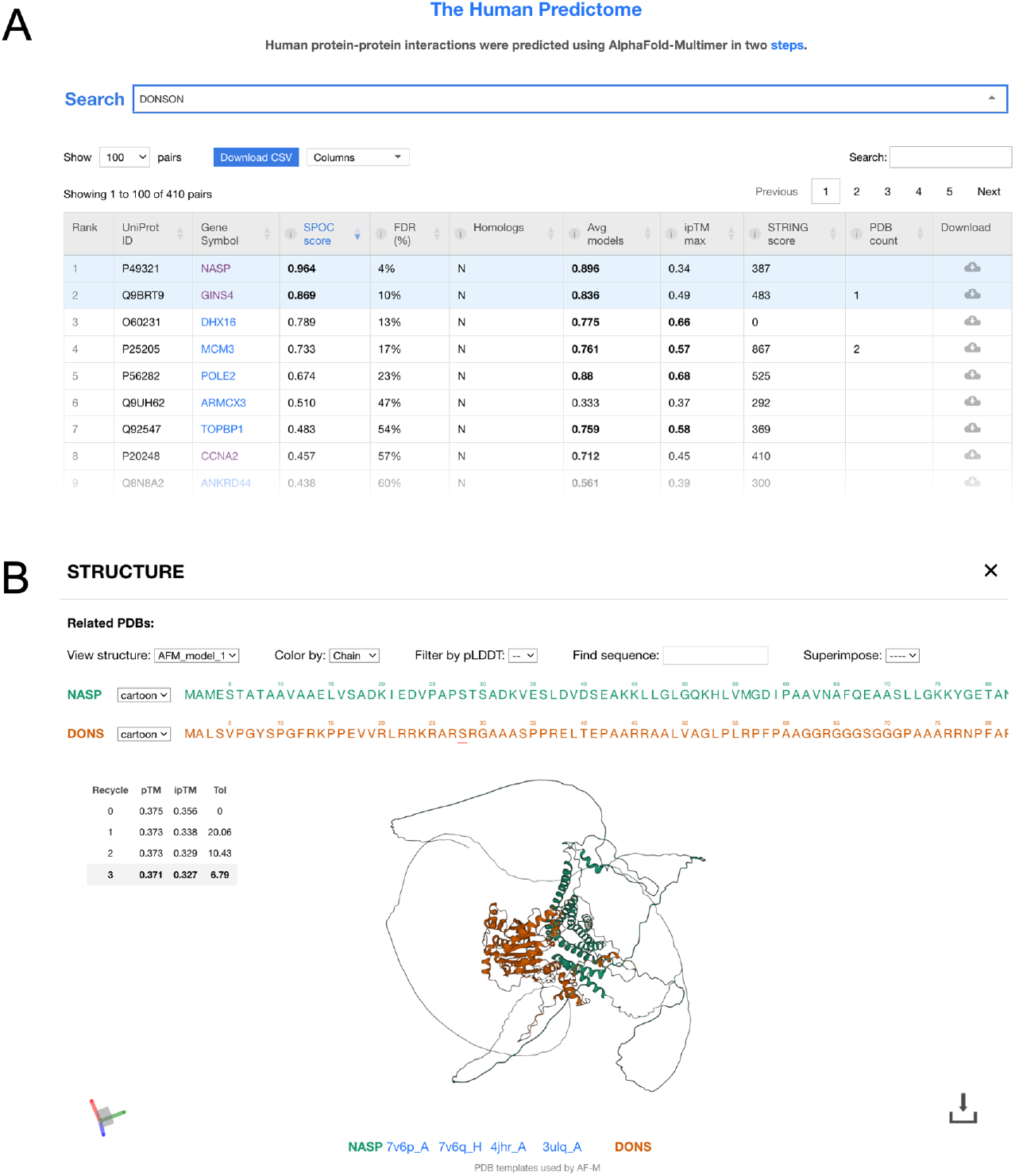
A web portal for viewing the human predictome. Screenshot from the human predictome website displaying the table of hits for DONSON. **B**. Screenshot from the interactive structure viewer that appears upon selecting NASP in the table described in (A). The viewer allows users to superimpose homologous interactions from the PDB for comparative analysis.

### Hypothesis generation

The following examples illustrate how the predictome seeds new mechanistic hypotheses. The first case concerns the repair of DNA interstrand crosslinks (ICLs). When replisomes converge on an ICL, mono-ubiquitylated FANCI–FANCD2 clamps onto DNA near the damage and promotes recruitment of the nuclease XPF in complex with the scaffolding protein SLX4 (refs.^22–25^)(**Figure 4A**). However, the mechanism by which FANCI–FANCD2–Ub recruits XPF remains unknown. Our *in silico* screening identified a highly conserved motif near the N-terminus of SLX4 that is predicted to interact with the outer surface of FANCI in a configuration that is compatible with the FANCI–FANCD2–Ub complex structure (**Figure 4A**, pink arrowhead, **Figure 4B**, and **Figure S4A-B** for PAE plots; SPOC=0.65, FDR=25%). The limited interaction surface suggests that stable interaction might require additional contacts. Notably, SLX4 contains a ubiquitin-binding (UBZ) domain required for efficient repair^25,26^. Thus, we hypothesize that SLX4 is a coincidence sensor whose recruitment requires recognition of ubiquitin attached to FANCD2 and a specific surface on FANCI (**Figure 4A**, pink and blue arrowheads). Although FANCI and SLX4 have been functionally linked (STRING = 967), a direct interaction between these proteins has not previously been proposed.

**Figure 4.**
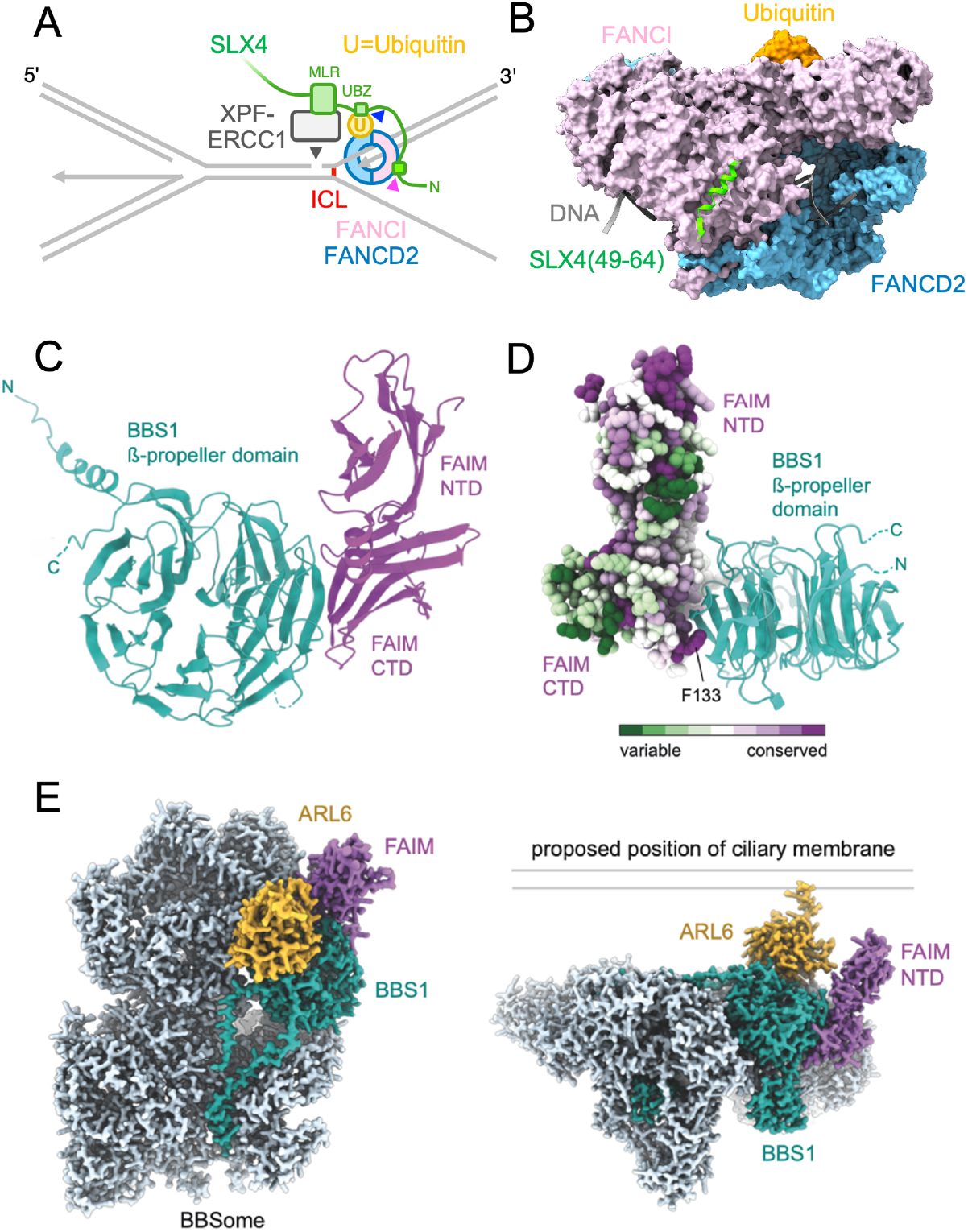
Hypotheses seeded by predicted interactions. **A**. Schematic model of interstrand crosslink (ICL) repair. Ubiquitylated FANCI–FANCD2 recruits the SLX4–XPF–ERCC1 complex. We propose that the UBZ and N-terminal domains of SLX4 contact Ubiquitin and FANCI (blue and pink arrowheads, respectively). This model requires that ubiquitin be reoriented from its pose in PDB:6VAF so that SLX4’s UBZ domain can bind the ubiquitin hydrophobic patch. **B**. The FANCI– SLX4(residues 49-64) prediction aligned to the FANCI–FANCD2– Ubiquitin–DNA structure (PDB:6VAF), shown in surface representation. **C**. Cartoon representation of the predicted interaction between the BBS1 β-propeller domain (cyan) and the C-terminal domain (CTD) of FAIM (purple). **D**. Residue conservation analysis mapped onto the atomic model of FAIM of the BBS1–FAIM complex. Highly conserved residues are shown in purple, and variable residues in green. The invariant F133 residue of FAIM protrudes into a hydrophobic cleft of BBS1. The N-terminal domain (NTD) of FAIM has a conserved patch of surface-exposed hydrophobic residues. **E**. Two views of a predicted BBSome– ARL6–FAIM complex. Left, BBSome (light blue), with BBS1 (cyan), FAIM (purple), and ARL6 (gold). Right, Complex orientated relative to a proposed position of the ciliary membrane.

Our screen also predicted associations between proteins not previously implicated in the same biological pathway. For example, we identified an interaction between Bardet-Biedl syndrome 1 (BBS1) and Fas apoptosis inhibitory molecule (FAIM) (**Figure 4C and Figure S4C;** SPOC=0.678, FDR=21%). BBS1 is a subunit of the octameric BBSome complex, which facilitates trafficking of transmembrane proteins within cilia^27–29^, whereas FAIM is an anti-apoptotic protein^30^. AF-M predicts an interface between the C-terminal domain of FAIM and the N-terminal β-propeller domain of BBS1 (**Figure 4C**). The interaction involves conserved surface patches on both proteins and buries an invariant and otherwise energetically unfavorable surface-exposed phenylalanine (F133) of FAIM^31^ (**Figure 4D**). Further confidence in the interaction comes from the absence of FAIM-like domains in any other human protein, the compatibility of the prediction with cryo-EM structures of the BBSome and BBSome–ARL6 complexes^32,33^ (**Figure 4E**), and the ability of AlphaFold3 to predict the entire BBSome–FAIM interaction with high confidence (**Figure 4E** and **S4D**). If FAIM binds the BBSome concomitantly with ARL6, which recruits the BBSome to the ciliary membrane, the N-terminal domain of FAIM would be positioned near the ciliary membrane (**Figure 4E**), positioning a conserved surface (**Figure 4D**) for interaction with a yet unidentified factor. Furthermore, mice deficient in BBS1 or FAIM exhibit similar phenotypes characterized by obesity and retinal degeneration^34–37^. Together, these observations raise the intriguing possibility of a pathway linking ciliopathy mechanisms to cell survival regulation. Future experiments are needed to confirm this interaction and explore its physiological role.

### Interpreting cryo-EM density maps

For low-resolution cryo-EM reconstructions, such as those derived from cryotomography, traditional density-guided methods^38,39^ often fail to unambiguously identify constituent proteins. To explore the utility of *in silico* PPI screening for interpreting subtomogram averages, we examined a recently published *in situ* structure of the mouse sperm central apparatus (CA)^40^. The CA comprises a pair of microtubules with interconnected complexes called “projections” that forms the center of the axoneme in sperm and motile cilia and that helps regulate motility. The subtomogram average revealed unassigned densities, particularly in the distal regions of the C2a and C1b projections, where local resolution is lowest (**Figure 5A**, yellow density). To help elucidate their composition, we assessed predicted interactions among proteins previously assigned to these projections (**Figure 5B-C, Figure S5 and Table S6**).

**Figure 5.**
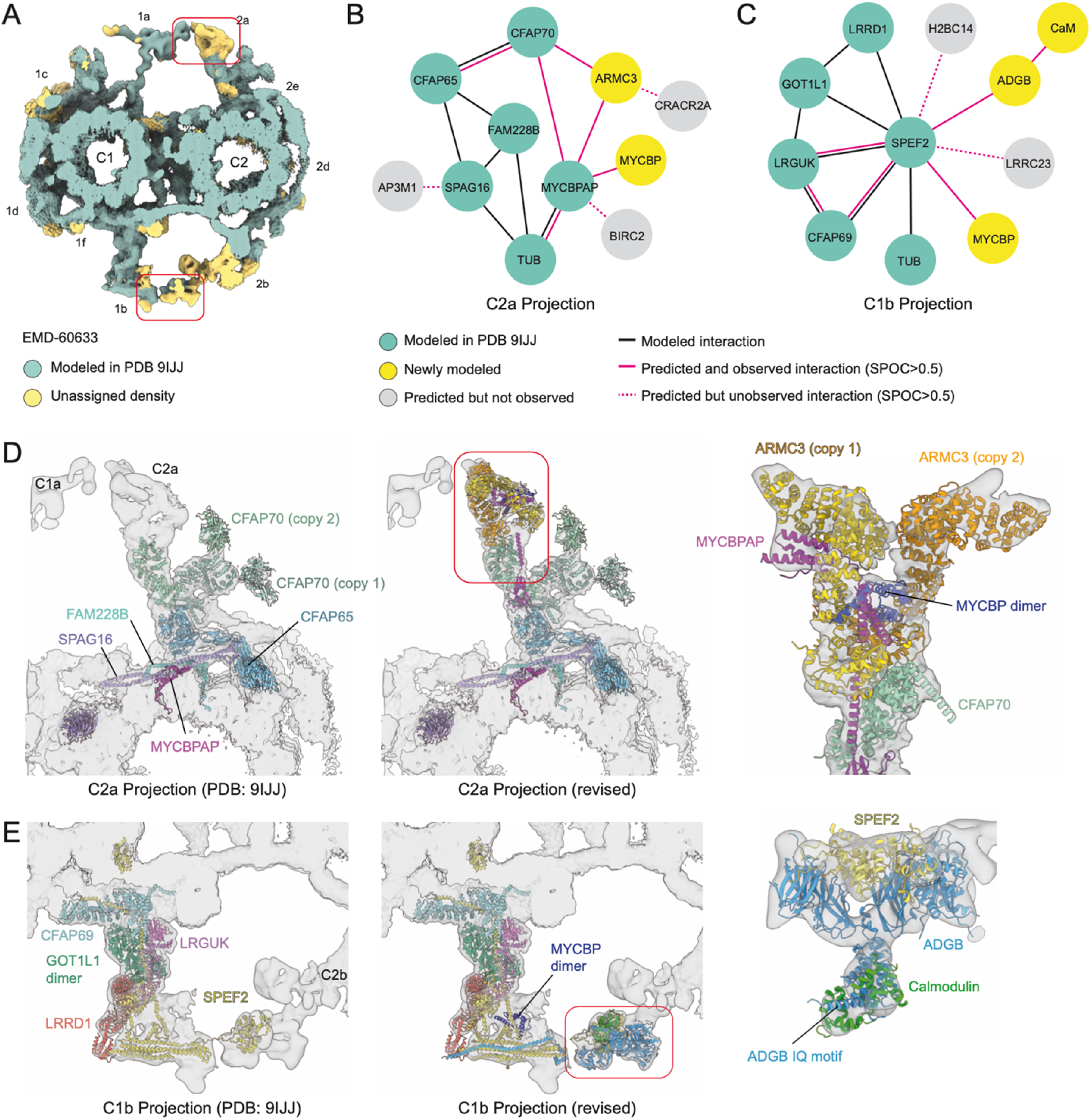
Improving atomic model with predicted PPIs. **A**. Isosurface rendering of the subtomogram average of the mouse sperm central apparatus (EMDB: EMD-60633) colored by whether it had been modeled in PDB 9IJJ, with yellow indicating unmodeled densities. C1 and C2 microtubule projections are labeled following convention; boxed areas show the distal regions of analyzed projections. **B-C**. Protein-protein interaction networks for the mouse sperm C2a (B) and C1b (C) projections, generated by integrating experimentally determined interactions from PDB 9IJJ with predicted interactions. Proteins in grey are predicted but not observed in the density. Calmodulin (CaM) was predicted by the known association with ADGB^77^. **D**. Left, Atomic model of the C2a projection from PDB 9IJJ, with subtomogram average density (EMD-60633) as a transparent isosurface. Middle, C2a model after incorporating predicted interactions. Right, zoom-in showing the fit of two copies of ARMC3 to the distal C1b region. One copy of ARMC3 interacts with MYCBPAP and a MYCBP dimer. **E**. Left, Atomic model of the C1b projection from PDB 9IJJ with subtomogram average density (EMD-60633) as a transparent isosurface. Middle, C1b model after incorporating predicted interactions. Right, zoom-in showing the fit of the SPEF2-ADGB-Calmodulin complex to the C1b projection density.

For the C2a projection, our analysis revealed an unmodeled interaction between the established subunits CFAP70 and MYCBPAP (SPOC=0.756, FDR=15%). We also detected two further subunits: MYCBP, which interacts with MYCBPAP (SPOC=0.65, FDR=25%), and ARMC3, which interacts with both CFAP70 (SPOC=0.542, FDR=41%) and MYCBPAP (SPOC=0.653, FDR=25%).

Although the MYCBPAP–MYCBP interaction was anticipated^41^, ARMC3 had not been previously recognized as a CA subunit. These predictions enabled construction of a more complete atomic model of the C2a projection (**Figure 5D; Table S7; Movie S1**). Two copies of ARMC3 are present per projection, with one bound by MYCBPAP and a MYCBP dimer, and the other making more extensive interactions with CFAP70. The interaction between ARMC3 and MYCBPAP, as well as the self-association of ARMC3, are supported by immunoprecipitation experiments^42,43^. The assignment of ARMC3 as a CA subunit is further reinforced by data showing that male *Armc3*-deficient mice are infertile with nearly immotile sperm^42^ and the association of human *ARMC3* variants with Multiple Morphological Abnormalities of the Flagella (MMAF), characterized by CA loss and axonemal disorganization^42,44^.

For the C1b projection, our *in silico* screen and subsequent modeling revealed that SPEF2 interacts with two previously unmodelled proteins, MYCBP (SPOC=0.683, FDR=21%) and ADGB (SPOC=0.843, FDR=11%) (**Figure 5C, E; Table S6; Movie S1**). Although ADGB had been assigned to the CA of *Tetrahymena thermophila* based on biochemical evidence^45^, its structure and location within the mammalian CA were unknown. Our model positions ADGB in the distal C1b projection together with SPEF2 and Calmodulin, a known interaction partner^46^ (**Figure 5E; Movie S1**). Genetic variants in human *ADGB* have been linked to male infertility and sperm axonemal disruption^46–48^. Although *ARMC3* and *ADGB* are both highly expressed in sperm, and their genetic loss manifests as male infertility, single-cell RNA-sequencing demonstrates that both genes are also expressed in multiciliated respiratory epithelial cells^49^ (**Figure S6**), indicating that their gene products are likely conserved CA subunits across mammalian motile cilia.

In conclusion, integrating *in silico* PPI screening with experimental structural data allows for the identification of subunits, clarifies the molecular organization of complex assemblies, and provides a framework for interpreting genetic variations associated with human disease.

### Validation of predicted PPIs

To explore a newly predicted PPI in detail, we focused on the replication factor DONSON. DONSON promotes assembly of the replicative CMG helicase by interacting with TOPBP1, GINS, and MCM3 (**Figure 6A**). It also makes a non-essential interaction with DNA Pol ε. While these known partners ranked in the top seven DONSON hits in our proteome-wide screen (**Figure 3A**), the most confident hit was the protein NASP (SPOC=0.96; FDR=4%). NASP maintains histone levels and binds with ASF1B to the histone H3-H4 heterodimer^50^. ASF1B in turn has been implicated in the incorporation of newly synthesized H3-H4 into daughter DNA strands by CAF-1^51^. AF-M predicts that two short, conserved segments in the N-terminal region of DONSON wrap around NASP (**Figure 6A-C**), which would preclude DONSON binding to Pol ε but not to GINS, TOPBP1, or MCM3 (**Figure 6A** and ref. ^6^). Importantly, DONSON and H3-H4 binding to NASP are predicted to be sterically incompatible (**Figure S7A**).

**Figure 6.**
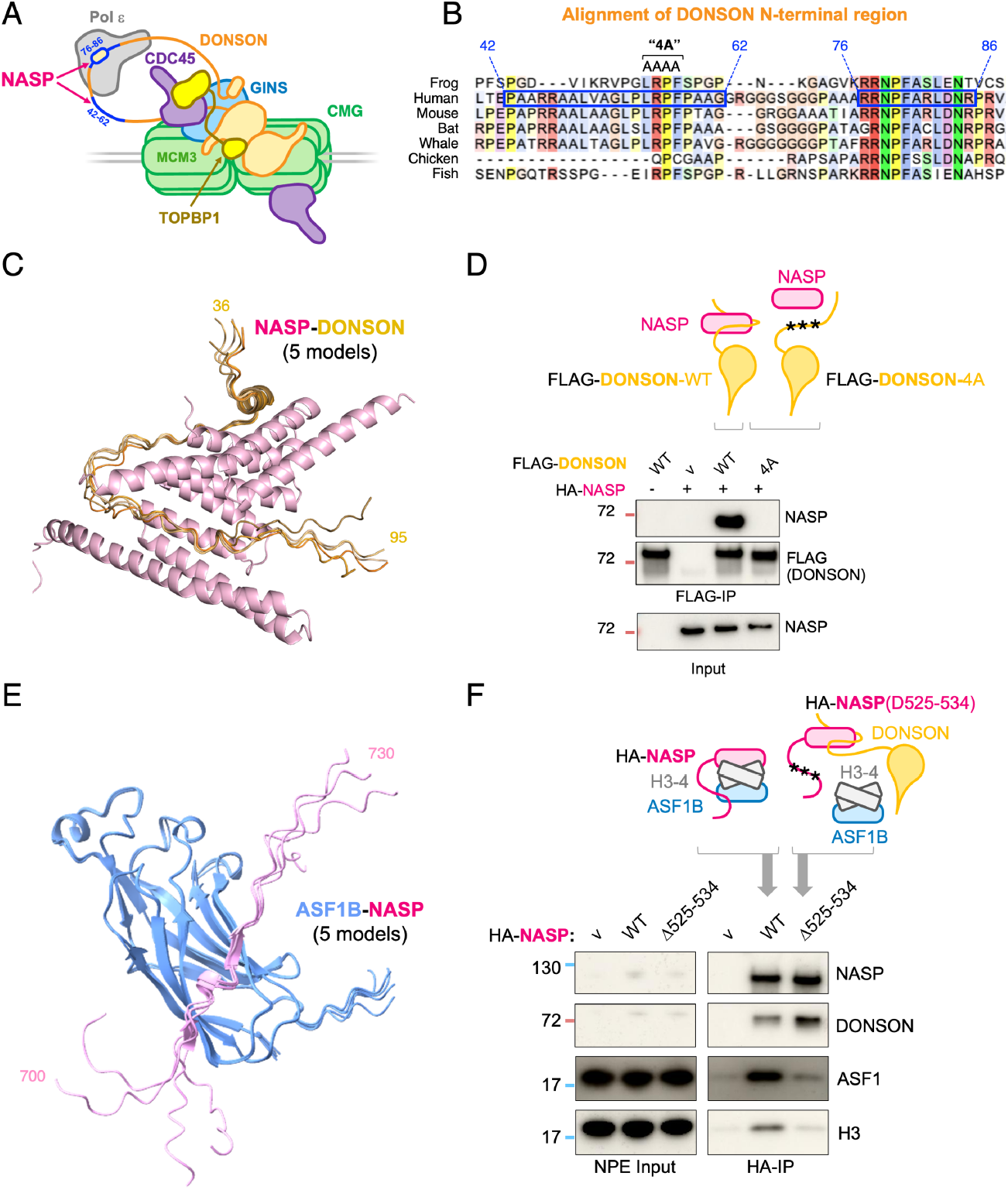
NASP interacts with DONSON and ASF1B. **A**. Schematic model of DONSON’s interactions. DONSON promotes CMG assembly by interacting with GINS (GINS4 subunit), Pol ε (POLE2 subunit), TOPBP1, MCM3, and DONSON (homodimerization). The second DONSON is thought to mediate similar interactions (not shown). The two short motifs in the N-terminal region of DONSON predicted to interact with NASP are indicated in blue, including approximate residue numbers. **B**. Sequence alignment of the N-terminal region of DONSON across seven species. The bipartite NASP-binding motif is indicated by the two blue boxes on human DONSON (same numbering as in (A)), and residues mutated to alanine to generate the “4A” mutant are indicated above the alignment. **C**. The five AF-M predictions of DONSON (residues 36-9 5) with NASP, superimposed by aligning NASP models. **D**. The N-terminal region of DONSON interacts with NASP, as predicted by AF-M. *Top:* schematic representation of DONSON– NASP co-IP results. *Bottom:* human HA-NASP and FLAG–DONSON or FLAG-DONSON^4A^ (where four amino acids predicted to interact with NASP were mutated) were co-expressed in wheat germ extracts, FLAG-DONSON was precipitated, and the indicated proteins were visualized by immunoblotting. The lower gel shows the input. **E**. The five AF-M predictions of human NASP (residues 700-730) with ASF1B, superimposed by aligning ASF1B models. **F**. The C-terminal disordered region of *Xenopus* NASP interacts with *Xenopus* ASF1B, as predicted by AF-M (the interaction is very similar to that of human NASP and ASF1B shown in (C), but Xenopus NASP is smaller so the ASF1B-binding motif is in a different location). *Top:* schematic representation of the NASP–ASF1B co-IP results. *Bottom:* HA–NASP or a NASP mutant carrying the indicated deletion (Δ525–534) was added to NPE, recovered, and analyzed by immunoblotting.

To address whether DONSON interacts with NASP, we used *in vitro* translation to express human NASP with wild type human DONSON or with a DONSON mutant in which four amino acids (LRPF) predicted to interact with NASP were mutated to alanine residues (DONSON^4A^; **Figure 6B**). DONSON^WT^ but not DONSON^4A^ co-precipitated NASP (**Figure 6D** and **S7B**). Conversely, in mammalian cells, NASP mutants designed to disrupt histone binding failed to interact with DONSON^52^, as expected if H3-H4 and DONSON bind the same surface on NASP (**Figure S7A**). Both DONSON^WT^ and DONSON^4A^ supported DNA replication in frog egg extracts, indicating that the DONSON–NASP interaction is not required for replication initiation, as predicted (**Figure S7C**). Interestingly, acute depletion of DONSON from human cells causes an abrupt cessation of DNA synthesis, suggesting a role in replication elongation^6^. A similar phenotype is observed in human cells depleted of CAF-1 or NASP^53,54^, consistent with the possibility that the DONSON–NASP interaction contributes to chromatin assembly at the replication fork. There was no defect in supercoiling of newly replicated DNA when we disrupted the DONSON–NASP complex in egg extracts (**Figure S7D**), suggesting that replication-coupled chromatin assembly was normal in this system. However, compared with mammalian cells, where free histones account for only ∼1% of the total histone pool^55^, egg extract contains a large excess of free histones, which might mask the requirement for the DONSON-NASP interaction in chromatin assembly.

NASP and ASF1B were previously shown to interact indirectly by binding to the same H3-H4 dimer^50^. Interestingly, NASP’s C-terminal region was predicted with high confidence to engage in a previously unrecognized interaction with ASF1B (**Figure 6E**; SPOC=0.915; FDR=7%). Accordingly, NASP co-precipitated ASF1B from frog egg extracts, but not when the ASF1B-binding motif in NASP was mutated (**Figure 6F**). Loss of the NASP-ASF1B interaction had two additional effects: it reduced co-precipitation of histone H3 with NASP, suggesting cooperative binding of NASP and ASF1B to the H3-H4 dimer (**Figure 6F**); it also increased the recovery of DONSON with NASP, as expected if DONSON competes with H3-H4 for NASP binding (**Figure 6F** and **Figure S7E**). Based on these observations, we speculate that DONSON recruits ASF1B–NASP–H3-H4 complexes to the replisome by binding NASP (**Figure S8**). This interaction releases H3-H4 from NASP, promoting H3-H4 transfer to ASF1B. Finally, ASF1B interacts with CAF-1^56^, which receives the transferred H3-H4 for deposition on DNA. Future experiments are necessary to directly test the functions of these newly identified DONSON–NASP and NASP– ASF1B interactions.

## Discussion

To identify new protein-protein interactions in the human proteome, we developed KIRC, a rapid classifier that allowed us to assess the interaction likelihood for all 200 M human protein pairs. The top 1.6 M pairs were then modeled with AlphaFold-Multimer and scored using SPOC. Our pipeline yielded ∼16,000 high-confidence PPIs, nearly a third of which were previously uncharacterized, representing a large expansion of structurally modeled human PPIs. To promote broad accessibility, this dataset is hosted at predictomes.org, along with analysis tools that allow rapid triage and hypothesis building.

Although our approach has identified many novel PPIs, it has limitations. Notably, at the thresholds applied (KIRC=0.088; SPOC=0.86), we estimate the recall of KIRC and SPOC to be ∼68% and ∼27%, respectively. In addition, using our protocol, AF-M recall is only ∼40% of known pairs in the PDB^14^. Combined, this would yield an overall recall for our pipeline of only ∼7% when applying a high-confidence threshold. Assuming our 16k set is associated with a 10% FDR, this translates to ∼14.4k true pairs, representing ∼7% of the ∼200k pairs thought to comprise the human interactome. In an independent estimate of recall, among 210 interactions deposited in the PDB after KIRC training (**Figure S2B**), ∼7% exceeded the SPOC threshold of 0.86 (**Table S8**), which is also similar to the recall in a recently reported computational interactome^2^. Relaxing the SPOC cutoff to 0.5 increased recovery to 113k pairs; with a corresponding increase to 50% FDR, this larger set is expected to contain ∼56k true pairs or ∼28% of the full 200k interactome. In summary, at high confidence, we likely recovered only a small fraction of the human interactome (7%), whereas at lower confidence, the yield is considerably higher (28%). As illustrated in the results section, many lower-scoring interactions generated compelling hypotheses that were validated using mutagenesis or by fitting to cryo-EM maps, demonstrating the utility of this larger set of predictions.

Most of the confident pairs we identified involve well-studied proteins. This bias likely reflects the fact that KIRC and SPOC were trained on biological omics data to reduce false positive interactions. As a result, proteins that are challenging to study or that show restricted developmental or tissue expression are penalized, perpetuating existing coverage limitations. One possibility for addressing this problem would be to fold all 200 M protein pairs and rely more heavily on structural features to score the predicted interactions. This approach, which should become feasible with more efficient structure prediction models, is expected to increase recall, but at the expense of a much higher FDR. Even with a higher FDR, it may be possible to use targeted experiments to identify which of the top structural hits are *bona fide* interactors. PPIs that differ fundamentally from the PDB structures used to train AF-M might still be difficult to detect. Overcoming bias to illuminate the “dark interactome” will likely require a concerted effort using multiple approaches.

Although large-scale experimental validation of the predicted PPIs is not possible, we and others have previously confirmed numerous predictions with both modest and high SPOC scores^6,9–12,57^ (**Table S4**). Here, we additionally verified the predicted DONSON–NASP and NASP–ASF1B interactions by demonstrating that mutations in predicted interfaces disrupt the interactions. Some of the new predictions involve contacts between proteins previously known to operate in the same pathway (e.g. NASP–ASF1B and FANCI–SLX4), whereas others involve proteins thought to operate in different processes (e.g. DONSON–NASP and BBS1–FAIM). Many of the predictions lead to compelling and testable hypotheses, such as the model that SLX4 is recruited to ICLs via a direct interaction with FANCI, or the notion that the BBS1– FAIM interaction links cilia dysregulation to cell death. In another powerful application, the predicted PPIs provided plausible interpretations of low-resolution experimental density maps where traditional density-guided methods failed.

In summary, by uncovering thousands of new, high confidence PPIs, the predictome is expected to seed novel mechanistic hypotheses across the human proteome and thereby accelerate progress towards a more complete understanding of cell physiology.

## Materials and Methods

### Assembling pair datasets for classifier training and testing

For KIRC training and evaluation, we assembled two sets of positive pairs and four sets of negative pairs. The two positive sets consisted of: (1) RefSet^PDB^ (N = 8,924), all unique human protein pairs that directly interact in the PDB as of 2024-04; and (2) RefSet^XLMS^ (N = 2,684, from^14^ but without applying a contact positive criterion), pairs captured in one of 20 large-scale crosslinking mass spectrometry (XLMS) experiments, where at least one of three AF-M structure predictions is geometrically consistent with the reported crosslinks (inter-residue Cα distance < 36 Å). For the negative sets, we generated RefSet^PDB_Decoy^ (N = 25,202), a set composed of all unique combinations of protein pairs found in the same complex or structure but not directly interacting. RefSet^Random^ (N = 204,151) included purely random interactions generated by randomly pairing proteins from the human proteome. This same random pairing approach was applied to the XLMS proteins to generate RefSet^XLMS_Random^ (N = 51,614). Finally, RefSet^XLMS_Decoy^ (N = 6,001) included all pairs captured in the XLMS experiments where none of the three AF-M predictions yielded a structure consistent with the reported crosslinks. All pairs are listed in **Table S2**.

### AlphaMissense data processing

Precomputed amino acid substitution scores for the human proteome were downloaded from AlphaMissense^58^: https://console.cloud.google.com/storage/browser/dm_alphamissense;tab=objects?pli=1&prefix=&forceOnObjectsSortingFiltering=false. For each residue, the AlphaMissense score was averaged across all 19 possible missense variants to produce a single, per-residue value. These values were then loaded into a JSON dictionary where each key is a UniProt ID that points to a numeric vector (with a length equivalent to the amino acid count of the protein) where each entry is a number from 0 to 100 that represents the averaged missense score predicted for mutating the residue at the corresponding position to a different amino acid.

### RNA co-expression data

Human mRNA co-expression profiles were downloaded from coexpressDB^59^: https://zenodo.org/record/6861444/files/Hsa-u.v22-05.G16651-S245698.combat_pca.subagging.z.d.zip. For each gene, co-expression scores were ranked (high to low) and the top 500 pairs retained. ENTREZ gene IDs were then mapped to canonical UniProt entry names using the UniProt mapping tool. Pairs that failed the mapping process failed were discarded. Score values were used as supplied by the database.

### DEPMAP data

Gene effect data from CRISPR knockout screens were downloaded from DEPMAP^60^: https://depmap.org/portal/download/custom/. Every protein was converted into a DEPMAP vector of length n = 1,095 (n = # profiled cell lines) where every entry/dimension in the vector corresponds to the Chronos output (gene effect) for that gene in a cell line.

### BioGRID Open Repository of CRISPR Screens (ORCS) data processing

CRISPR knockout screen data for human cell lines were downloaded from BioGRID ORCS^61^ : https://downloads.thebiogrid.org/File/BioGRID-ORCS/Release-Archive/BIOGRID-ORCS-1.1.15/BIOGRID-ORCS-ALL-homo_sapiens-1.1.15.screens.tar.gz. Each gene was mapped to a canonical UniProt ID. For each gene, its appearance across all the CRISPR screens was converted into binary vectors of length n = 1,243 (n = # of screens) where each index represents whether that gene was considered a “hit” (0 = no hit, 1 = hit) by the criterion employed by a specific screen.

### BioGRID interaction data

Interaction data were downloaded from BioGRID release 4.4.225: https://downloads.thebiogrid.org/File/BioGRID/Release-Archive/BIOGRID-4.4.225/BIOGRID-ALL-4.4.225.mitab.zip (August 2023) and filtered to retain only human proteins (taxid:9606). Human protein pairs were then identified using UniProt IDs included in the file. The number of times a unique human pair was found in this file was then used as the biogrid_detect_count feature and encoded into a nested dictionary JSON file where UniProt IDs are used as keys and point to the detect count value.

### Protein localization predictions

Protein localization probabilities were predicted for all canonical Swiss-Prot reviewed sequences for the human proteome downloaded from UniProt using a local installation of DeepLoc 2.0 (ref.^62^): https://services.healthtech.dtu.dk/cgi-bin/sw_request?software=deeploc&version=2.0&packageversion=2.0&platform=All. The ESDM1B “fast” model was used to predict localization probabilities for 10 different compartments per protein. These values were then loaded and stored in a JSON dictionary where each key is a UniProt ID that points to a numeric vector with the 10 localization probabilities output by DeepLoc 2.0.

### H5 protein embeddings

Per-protein embeddings (vectors of length 1,024) were retrieved for all reviewed UniProtKB Swiss-Prot human entries from UniProt: https://ftp.UniProt.org/pub/databases/UniProt/current_release/knowledgebase/embeddings/UP000005640_9606/per-protein.h5^63^.

### STRINGDB scores

Human protein-protein association scores were obtained from STRING v12 (ref.^16^): https://stringdb-downloads.org/download/protein.links.detailed.v12.0/9606.protein.links.detailed.v12.0.txt.gz. Each entry in the file lists a pair of proteins identified by their STRINGDB ID consisting of the taxon ID (9606 for humans) concatenated with an ENSEMBL protein id. These ENSEMBL protein ids were mapped to UniProt IDs using UniProt’s mapping API. In cases where this mapping yielded non-canonical UniProt IDs or non-SwissProt entries, these ENSEMBL protein ids were mapped to genes and then each gene was mapped to the canonical SwissProt UniProt ID.

### Co-fractionation mass spectrometry data analysis

Aggregated human co-fractionation mass spectrometry data^64^ were downloaded from Zenodo: https://zenodo.org/records/8005773/files/Homo_sapiens.tar?download=1. We extracted data from the metric=cosine-missing=noise-transform=none-normalize=none.tsv file by examining every row (protein-pair) and calculating the row mean, median, and max values across all columns (experiments). Missing values were ignored.

### FoldSeek cluster data retrieval

Precomputed FoldSeek cluster data^65^ for all canonical human proteins were downloaded from AlphaFoldDB^66^ REST API: https://www.alphafold.ebi.ac.uk/api/cluster/members/{uniprot_id}?cluster_flag=AFDB%2FFoldseek&records=5000&start=0&sort_direction=DESC&sort_column=averagePlddt. For each protein we recorded the number of proteins in the cluster, their average pLDDT values, and their residue lengths. These were then stored and subsequently used to generate pairwise features.

### ProteomeHD correlation data

ProteomeHD correlation data^67^ were downloaded from: https://www.proteomehd.net/download_file/S1. The CSV file was converted into a JSON format where each protein is encoded as a key (UniProt ID) that points to a numeric vector of length 294 with floating point values representing SILAC ratios recording protein abundance changes in response to 294 biological perturbations/experiments. If no value was present in the CSV file, we recorded a 0 for that position.

### PaxDb data processing

Data for human protein abundances as aggregated across mass spectrometry experiments was downloaded from PaxDB^68^: https://pax-db.org/downloads/5.0/datasets/9606/9606-WHOLE_ORGANISM-integrated.txt. The STRINGDB_IDs were mapped to canonical UniProt IDs, and the data was converted into a JSON representation where each protein is encoded as a key (UniProt ID) that points to a float value (the proteomic abundance). This same procedure was used to create a second JSON data file (multi_sample) where each protein points to a vector representing abundances for three different PaxDb datasets with at least 80% proteome coverage: (https://pax-db.org/downloads/5.0/datasets/9606/9606-GPM_2012_09_Homo_sapiens_-_ensembl.txt, https://pax-db.org/downloads/5.0/datasets/9606/9606-LIVER-integrated.txt, https://pax-db.org/downloads/5.0/datasets/9606/9606-TESTIS-integrated.txt)

### DisGeneNet similarity cosine data

Cosine similarities of DisGeNET^69^ gene phenotype association scores for all gene-gene pairs were downloaded as a matrix from Harmonizome version 3.0 (ref.^70^): https://maayanlab.cloud/static/hdfs/harmonizome/data/disgenetdisease/gene_similarity_matrix_cosine.txt.gz. The floating-point scores were converted to integers by multiplying them by 1,000 and rounding to the nearest integer. Gene names were then mapped to UniProt IDs and all matrix data was converted into a JSON format where each UniProt ID is a key that points to a nested object which contains secondary UniProt IDs as keys that map to the final cosine scores.

### Training the Knowledge Informed Rapid Classifier (KIRC)

As with SPOC, we developed KIRC using a random forest–based machine learning framework (scikit-learn). This approach was selected for its rapid training speed, interpretability, strong performance on tabular datasets, and inherent resistance to overfitting. All possible combinations of datasets (2 positive, 4 negative; **Table S2**) were used to train random forest classifiers, except for any combinations lacking positive or negative pairs. Each dataset was represented by fractional subsets corresponding to 0, 0.33, 0.66, or 1 of its total size to generate the full range of training combinations. This resulted in a total of (2^4^ – 1) × (4^4^ – 1) = 3,825 dataset combinations. Each of the classifiers was instantiated as an instance of the scikit-learn RandomForestClassifier class with the following command: RandomForestClassifier(n_estimators=250, random_state=17232, criterion=‘log_loss’, bootstrap=False). Classifiers trained on 100% of the positive sets (RefSet^XLMS^ and RefSet^PDB^) were excluded to retain positives for testing. Any classifier that did not use decoys during training were also excluded to select classifiers that are likely better suited to handle this particularly challenging class of pairs. Finally, we retained only the subset of classifiers trained on some proportion of positive pairs derived from both the PDB and XLMS to ensure thorough representation of positive, ground truth interactions. After these pruning steps, 1,921 classifiers remained, which were evaluated on their ability to rank 39 test pairs excluded from the training sets.

The classifiers were sorted by two criteria (**Table S3**): the median rank (ascending), followed by the number of target pairs assigned a rank below 100 since we anticipated being able to fold approximately 100 pairs per human protein on average (descending). Post sorting, three classifiers achieved equivalent performance with median rank 8 and 31 pairs with rank below 100. Two of the three final classifiers trained on 0% RefSet^PDB_Decoy^, while one classifier trained on RefSet^PDB_Decoy^ but 0% RefSet^XLMS_Decoy^. Given that the RefSet^PDB_Decoy^ is based on decoys from the highest reliability dataset (PDB), we selected the classifier that was trained on these pairs as the final KIRC model. KIRC was trained on a dataset with 33% of the XLMS positive pairs, 67% of PDB positive pairs, 0% of RefSet^XLMS_Decoy^ pairs, 33% of RefSet^PDB_Decoy^ pairs, 67% of RefSet^XLMS_Random^ pairs, and 100% of RefSet^Random^ pairs.

### AlphaFold-Multimer (AF-M)

We used a modified version of ColabFold v1.5.2 to run AF-M. All our predictions used AF-M version 3 weights, models 1, 2, and 4 with 3 recycles, templates enabled, 1 ensemble, no dropout, and no AMBER relaxation. The Multiple Sequence Alignments (MSAs) supplied to AF-M were generated by a local installation of MMseqs2 operating with default settings. All MSAs consisted of vertically concatenated paired and unpaired MSAs for the query proteins. Predictions were run on 80 GB A100 NVIDIA GPUs. Given the memory limitations of these GPUs, we only ran sequences where the total combined length was less than 3,600 amino acids.

### PDB interacting homolog pair identification

To determine how many structurally resolved protein pairs in the PDB correspond to a homologous human pair, all PDB sequences were downloaded as a text file on April 16, 2025. Non-protein sequences were removed, resulting in a FASTA file containing only protein chain data. Protein chains containing fewer than 8 amino acids were excluded to eliminate short peptides. The remaining sequences were mapped to the canonical human proteome, obtained from UniProt, using MMseqs2 with the following command: mmseqs easy-search 20250416_pdb_seqres_proteins_gt7aa.fasta hs_proteome_uniprot_id_map.fasta 20250416_pdb_uniprot_aln.tsv tmp_dir -s 7.5 --min-aln-len 6 -- min-seq-id 0.3 -e 0.1 --exhaustive-search --format-mode 4. This search was configured for high sensitivity and required a minimum alignment of 6 residues and ≥30% sequence identity. For each unique PDB chain, only the best alignment was retained, prioritizing (1) highest percent identity, (2) lowest E-value, and (3) longest alignment.

Next, we downloaded all released PDB models. Using our previously generated mapping of PDB chains to human protein chains, we identified cases where two or more chains co-occurred in the same structure. We then assessed whether any of these human mappable chains were in contact, defining contact as ≥10 interchain residue pairs with any atom–atom distance <5 Å. This analysis yielded 269,072 interacting chain pairs, corresponding to 14,431 unique human protein pairs with a solved homologous interaction in the PDB.

To account for redundancy within the human proteome, we clustered the proteome using MMseqs2 at 90% identity and 90% coverage with the following command: mmseqs easy-cluster hs_proteome_uniprot_id_map.fasta hs_clusters tmp_dir - -min-seq-id 0.9 -c 0.9. Post clustering, we expanded the interaction list by including all pairwise combinations of proteins from any two clusters that contained interacting members (**Figure S2A**). The resulting set of 24,755 protein pairs was treated as the final list of PDB-supported (PDB+) human binary interactions for the majority of downstream analyses. For the FDR analysis outlined below, we used a more inclusive PDB+ set generated by clustering the proteome at 50% identity and 70% coverage.

### SPOC false discovery rate (FDR) analysis

Positive pairs were first sourced from those excluded during KIRC training. Pairs previously used for SPOC training were removed, leaving 2,881 positives for this analysis. From these, 250 were randomly sampled. In parallel, 324,675 random pairs were generated by pairing canonical human proteins (1.3 × 999 × 250; a 30% excess was used over the required 999 × 250 negative pairs to ensure enough remained for testing after pruning). These random pairs were filtered to exclude any inadvertent positives, defined as pairs present in IntAct (evidence count ≥2 with sequence identity ≥50% and coverage ≥70%), in the PDB homology interaction set (identity ≥50% and coverage ≥70%), or with STRING scores >990 (identity ≥90% and coverage ≥90%). From the surviving random negative pairs, 249,750 pairs were sampled.

The final dataset thus contained 250 positive and 249,750 negative pairs at a 1:999 ratio, approximating the expected positive:negative ratio in the human proteome. KIRC and SPOC scores were then calculated for all pairs to determine true positive and true negative rates across different cutoffs, which were converted to estimated FDR values under the assumption of 200,000 true pairs and 199.8 million false pairs in the human proteome. This procedure was repeated 100 times independently to ensure robustness, and reported values represent the means across these 100 simulations.

### Post-run PDB validation analysis

To assess post-prediction performance, we first identified all human pairwise interactions that are reported in the PDB, including those inferred by homology. These structurally resolved interactions were then clustered via sequence homology. Pairs were grouped into the same cluster if the pairs consisted of proteins from the same sequence clusters (≥90% identity, ≥90% coverage). We next determined the earliest date at which any of the cluster members was resolved in a structure published in the PDB (**Figure S2B**). Based on this, 210 unique interaction clusters were first published after October 1, 2024 (KIRC data cutoff) and before April 16, 2025. All recall analysis was performed at the cluster level rather than for individual pairs within clusters. For each cluster, the highest SPOC score among its member pairs was used to determine whether the cluster was represented in the final 16k set.

### Gene Ontology (GO) term analysis

A list of GO terms associated with human proteins^71^ was downloaded from the GOA database: https://ftp.ebi.ac.uk/pub/databases/GO/goa/proteomes/https://ftp.ebi.ac.uk/pub/databases/GO/goa/proteomes/25.H_sapiens.go a (May 2025). A complete list of all GO terms and their definitions was similarly downloaded from https://purl.obolibrary.org/obo/go/go-basic.obo (July 2025). A custom Python script was used to parse these two files and produce a JSON file that maps each UniProt ID to an array of unique GO terms. GO terms were additionally collapsed into high similarity clusters by using the Levenstein text similarity distance to compare GO term description text with group inclusion requiring a normalized Levenstein similarity (1 – Levenstein(a, b)/max(len(a), len(b)) of > 0.75 to the founding cluster representative term.

For every protein pair, each protein was first assigned to a unique set of GO term clusters. Then, a Jaccard similarity score for GO terms was calculated by taking the intersection of GO term clusters divided by the union of the GO term clusters.

Gene set enrichment analysis^72^ was performed by using the online tool at https://geneontology.org/ and inputting a list of all proteins (UniProt IDs) identified in the 16k set against the baseline of the entire human proteome.

### Predictomes web server

The predictomes.org website is deployed as a scalable web platform on Amazon Web Services (AWS), extending the Structural Biology Cryo-EM Cloud Infrastructure recently developed for the SBGrid Consortium^73^. It utilizes Amazon CloudFront, a global content delivery network (CDN), to cache and deliver content with low latency to users worldwide. All requests are routed through an AWS Application Load Balancer (ALB), which distributes incoming traffic across multiple container instances and only forwards requests to healthy backend targets. The web application runs inside Docker containers managed by Amazon Elastic Container Service (ECS), AWS’s fully managed container orchestration service. This design ensures high availability and allows the service to automatically scale to handle increased demand.

The application’s data storage leverages AWS managed services for reliability. Analysis metadata is stored in a MySQL relational database hosted via Amazon Relational Database Service (RDS), which easily scales to accommodate demand. Large data files (such as predicted protein structures and other results) are stored via Amazon Simple Storage Service (S3). These files are distributed to users via CloudFront, which serves data from edge caches rather than the origin server, improving download speeds. The API (application programming interface) endpoints that deliver analysis data also support CloudFront caching, to minimize database load and speed up repeated queries. The platform runs compute-intensive analyses of user-uploaded data as separate tasks on the ECS cluster. Specifically, the AlphaFold 3 (AF3) analysis pipeline and the SPOC (Structure Prediction and Omics-informed Classifier) scoring routine execute as on-demand container tasks, isolating heavy computations from the main web server. This approach prevents long-running analyses from reducing site responsiveness. After processing, results are delivered to users via email using Amazon Simple Email Service (SES).

All infrastructure provisioning and configuration is managed through Terraform, an open-source “infrastructure as code” tool that allows the entire AWS environment to be defined and version-controlled in code. The deployment process also incorporates a CI/CD (Continuous Integration and Continuous Deployment) pipeline, which automatically tests code changes and rolls out updates to the production environment, ensuring that new features and fixes are released reliably and with minimal downtime.

### Atomic model building and refinement

The subtomogram average of the mouse sperm central apparatus was retrieved from the Electron Microscopy Data Bank (EMDB: EMD-60633) and the corresponding atomic model was obtained from the Protein Databank (PDB: 9IJJ). For this study, the C2a projection was defined as comprising FAM228B, CFAP65, CFAP70, SPAG16, and MYCBPAP, whereas the C1b projection was defined as comprising CFAP69, SPEF2, LRGUK, GOT1L1, and LRRD1. Protein-protein interaction predictions for each of these proteins were retrieved from predictomes.org and filtered by a SPOC score threshold greater than 0.5.

Because the initial predictions used human sequences and the structure was derived from mouse sperm, all models were re-predicted with mouse sequences using AlphaFold3 (ref.^74^). AlphaFold3 was also used to predict multisubunit complexes, combining pairwise interactions (**Figure S5B**). Predicted complexes were fitted to the subtomogram average density using ChimeraX^75^ and Coot^76^. Regions of the model outside the density and with low pLDDT scores were trimmed using Coot.

For the C2a projection, fitting revealed two copies of ARMC3: one associated with CFAP70, and a second bound to MYCBPAP and a MYCBP dimer. In the C1b projection, a MYCBP dimer also corresponded better to the density associated with SPEF2 than a MCYBP monomer. Additional density near ADGB was modeled as Calmodulin (UniProt: P0DP26), which has been shown experimentally to bind the IQ motif of ADGB^77^.

Atomic models were refined using real-space refinement in Phenix v.1.21.2-5419 (ref.^78^). A conservative overall resolution cutoff of 12 Å was set during refinement. This value is lower than resolution estimations for the C1b and C2a projections (8.9 and 7.8 Å, respectively)^40^ but more accurately represents the local resolution at the distal regions of the maps where the most substantial model modifications were introduced. Each model underwent two consecutive refinements runs, with each run consisting of five macrocycles and 100 iterations per macrocycle. Each refinement cycle included coordinate minimization, sidechain flips, and local grid search. Reference model restraints, secondary structure restraints, and Ramachandran restraints were applied throughout to maintain the good starting geometry of the input model. Model quality was assessed following refinement using MolProbity^79^. Refinement statistics are reported in **Table S7**.

Figures showing subtomogram averages and atomic models were generated using ChimeraX^75^.

### Protein Conservation Analysis

Residue conservation analysis for FAIM and BBS1 was performed using ConSurf^80^ with default parameters.

### FANCI–FANCD2–SLX4 modeling

Residues 1-100 of human SLX4 were folded with FANCI using AlphaFold2.3 (ColabFold). This prediction resembles the SLX4– FANCI complex predicted using full-length proteins in our *in-silico* screen. For **Figure 4B**, residues in SLX4 with a pLDDT score below 45 (residues 1-48 and 65-100) were hidden, and the complex was aligned to the cryo-EM-derived model of the ubiquitylated FANCI–FANCD2 complex (PDB:6VAF)^81^.

### Animal ethics

Egg extracts were prepared using female adult *Xenopus laevis* (Nasco Cat #LM0053MX). All experiments involving animals were approved by the Harvard Medical Area Standing Committee on Animals (HMA IACUC Study ID IS00000051-6, approved 10/23/2020, and IS00000051-9, approved 10/23/2023). The Harvard Medical School has an approved Animal Welfare Assurance (D16-00270) from the NIH Office of Laboratory Animal Welfare.

### DNA replication using egg extracts

*Xenopus* egg extracts and sperm chromatin were prepared as described^82^. To measure DNA replication efficiency and nucleosome assembly, replication licensing was carried out by adding plasmid DNA to the high-speed supernatant (HSS) of egg cytoplasm at a final concentration of 7.5 ng/µL. After 30 min, replication was initiated by mixing 2 volumes of nucleoplasmic extract (NPE) diluted 50% with 1xELB-sucrose (10 mM HEPES-KOH pH 7.7, 2.5 mM magnesium chloride, 50 mM potassium chloride, 250 mM sucrose) with 1 volume of the licensing reaction. For DONSON rescue experiments, 1 volume of immunodepleted NPE was supplemented with 0.1 volume of wheat germ extract expressing DONSON, or wheat germ extract containing an empty vector and preincubated at room temperature for 15 min prior to addition to the licensing reaction. At the indicated time points, samples of the replication reactions were quenched in 10 volumes of replication stop buffer (50 mM Tris-HCl pH 7.5, 25 mM EDTA, 0.5% SDS). Samples were treated with 4 µg of RNaseA for 30 min at 37 °C followed by 20 µg of Proteinase K (Roche 3115879001) digestion for 1 hour at 37 °C. Plasmid DNA was isolated by AMPure XP (SPRI beads, Beckman Coulter A63881) and the samples were then resolved on a native 1% agarose gel (supplemented with 1 µM chloroquine for analyzing supercoiling of newly replicated DNA). The dried gels were imaged on a Typhoon FLA 7000 PhosphorImager (GE Healthcare).

### Immunodepletions and rescue experiments in egg extracts

For immunodepletion of DONSON from egg extract, 0.5 volumes of 1 mg/mL affinity-purified DONSON antibodies^6^ were pre-incubated with 1 volume of Dynabeads Protein A (Invitrogen 10002D) by gently rotating at 4 °C overnight. 1.5 volumes of HSS or 70% NPE diluted with 1xELB were immunodepleted by three rounds of incubation with 1 volume of antibody-bound Dynabeads for 1 hour at 4 °C.

### Expression of proteins in wheat germ protein expression system

For protein expression, 3 volumes of TnT® SP6 High-Yield Wheat Germ Protein Expression System (Promega) were incubated with 2 volumes of 100 ng/μL pF3A WG (BYDV) Flexi vector (Promega) encoding human DONSON or NASP at 25 °C for 2 hours and used immediately. *Xenopus* DONSON or NASP were expressed in the same way.

### Immunoprecipitation

For immunoprecipitation, FLAG-DONSON and HA-NASP were expressed in Wheat Germ Protein Expression System. 0.15 volume of Pierce™ Anti-DYKDDDDK Magnetic Agarose (Thermo Scientific) was diluted with 1 volume of 2xELB-sucrose (120 mM HEPES-KOH pH 7.7, 5 mM magnesium chloride, 100 mM potassium chloride, 500 mM sucrose). 1 volume of Wheat Germ extract expressing FLAG-DONSON was incubated with 1 volume of magnetic agarose in 2xELB-sucrose for 1 hour at 4 °C. FLAG-DONSON-conjugated magnetic agarose was washed three times with 2xELB-sucrose, then 1 volume of Wheat Germ extract expressing HA-NASP and 1 volume of 2xELB-sucrose was added to the washed FLAG-DONSON-conjugated magnetic agarose. 0.5% of the mixture was taken as input. After 1 hour incubation at 4 °C, magnetic agarose was washed with 2xELB-sucrose three times. Agarose bound proteins were eluted by incubating with elution buffer (2xELB-sucrose with 0.25 mg/mL FLAG peptide) for 30 min at room temperature. For HA-NASP immunoprecipitations, Anti-HA Magnetic Beads (Pierce 88836) were used. The beads were pre-immobilized with HA-NASP proteins expressed in wheat germ extract and incubated with 30 µL 25% NPE for 1 hour at 4 °C. The beads were then washed three times with 2xELB-sucrose. Bound proteins were eluted by boiling each aliquot of beads with 1× Laemmli buffer.

### SDS-PAGE analysis and Western blotting in egg extracts

Protein samples were diluted with SDS sample buffer to a final concentration of 50 mM Tris pH 6.8, 2% SDS, 0.1% Bromophenol blue, 10% glycerol, and 5% β-mercaptoethanol and resolved on Mini-PROTEAN (Bio-Rad). Gels were then transferred to PVDF membranes (Thermo Scientific, PI88518). Membranes were blocked in 5% nonfat milk in 1x PBST for 1 hour at room temperature, then washed three times with 1x PBST, then incubated with primary antibodies diluted to 1:1000–1:5,000 in 1x PBST overnight at 4 °C. Following washes with 1x PBST three times, membranes were incubated for 1 hour at room temperature with light chain-specific mouse anti-rabbit antibodies (Jackson ImmunoResearch) at 1:10,000 dilution, or rabbit anti-mouse horseradish peroxidase-conjugated antibodies (Jackson ImmunoResearch) at 1:10,000 dilution in 5% nonfat milk in 1x PBST. Membranes were then washed three times with 1x PBST, developed with ProSignal® Pico ECL Spray (Genesee), and imaged using an Amersham ImageQuant 800 (Cytiva).

### Antibodies used for Western blotting

The following rabbit polyclonal antibodies were used for western blotting: DONSON (1:5,000 ref.^6^); *Xenopus* NASP (1:2,000; this study); human NASP (1:2,000; Invitrogen, PA5-55776); H3 (1:500; Cell Signaling, 9715S); ASF1B (1:1,000 ref.^83^). Monoclonal antibodies against HA (HA-Tag Rabbit mAb, Cell Signaling, 3724S) and FLAG (1:2,000; Monoclonal ANTI-FLAG® M2, Sigma Aldrich, F1804) were also used. Rabbit polyclonal antibodies against the C terminus of *Xenopus* NASP (Ac-CEESPLKDKDAKK-NH2) were prepared by BioSynth.

## Author Contributions

E.W.S. and J.C.W. conceived the project. E.W.S. performed all computation, except hyperparameter tuning and evaluation of KIRC, both of which were carried out by H.Z. H.Z. additionally created panels B-D in Figure 1 and panels C-D in Figure 2, as well as supplementary Tables S2-S5. The predictomes.org website and its analysis tools were designed and built by E.W.S with input from J.C.W. All experiments shown in Figures 6 and S7 were performed by E.R.; Y.L. first detected the interaction between NASP and DONSON. A.S. helped formulate a hypothesis regarding the FANCI-SLX4 prediction. A.B. formulated hypotheses involving ciliary proteins and built atomic models of the central apparatus projections. E.W.S., H.Z., A.B., and J.C.W. wrote the paper with feedback from the other authors.

## Acknowledgements

We thank NVIDIA corporation for use of a DGX server made available through the National Artificial Intelligence Research Resource (NAIRR) Pilot administered by the National Science Foundation (NAIRR). We thank F. Mattiroli, Anja Groth, and Michael Sun for helpful feedback. Deployment of predictomes onto AWS infrastructure was led by Ben Eisenbraun, Giorgos Boutsioukis, and Jason Key in the Sliz group at Harvard Medical School, leveraging their expertise in deploying scientific web applications. E.W.S. was supported by the National Science Foundation (DGE 2140743). A.B. was supported by NIH grant GM141109. A.S. was supported by NIH grant GM140400 and the G. Harold and Leila Y. Mathers Charitable Foundation. J.C.W. was supported by NIH grant HL098316 and the Dean’s Innovation Award. J.C.W. is an American Cancer Society research professor (RP-22-185-06-COUN) and a member of the Howard Hughes Medical Institute.

## Data availability

Atomic coordinates for the revised models of the C1b and C2a projections of the mouse sperm central apparatus have been deposited in the Protein Data Bank under accession codes 9YU4 (C1b projection; extended ID pdb_00009YU4) and 9YU3 (C2a projection; extended ID pdb_00009YU3).

## Table Legends

**Table S1: KIRC features**

Description of features used in the final KIRC interaction prediction model, along with their Gini importance scores.

**Table S2: KIRC dataset**

All protein pair datasets used to train and evaluate various interaction classifiers.

**Table S3: Classifier training results**

Performance of classifiers in ranking experiments across all classifier versions trained on various combinations of pairs from Table S2.

**Table S4: Ranking experiments**

List of positive interacting pairs from the literature that were used for ranking experiments during classifier training.

**Table S5: Novel pairs in the human predictome**

Table of metrics associated with high scoring protein pairs that were not identified in other interaction databases such as the PDB, STRING, and IntAct.

**Table S6: Cryo-EM density reanalysis**

Table of metrics associated with interactions that were fit into cryo-EM densities from the mouse sperm central apparatus complex (EMD-60633).

**Table S7. Refinement and validation statistics for the revised models of the C1b and C2a projections** Models were refined against EMD-60633.

**Table S8: Post KIRC PDB interaction analysis** Summary of metrics associated with proteins pairs that were first shown to directly interact in models deposited in the PDB after the finalization of KIRC.

**Movie S1: Remodeling of the mouse sperm central apparatus using interactions from the human predictome**.

---

**Figure S1:**
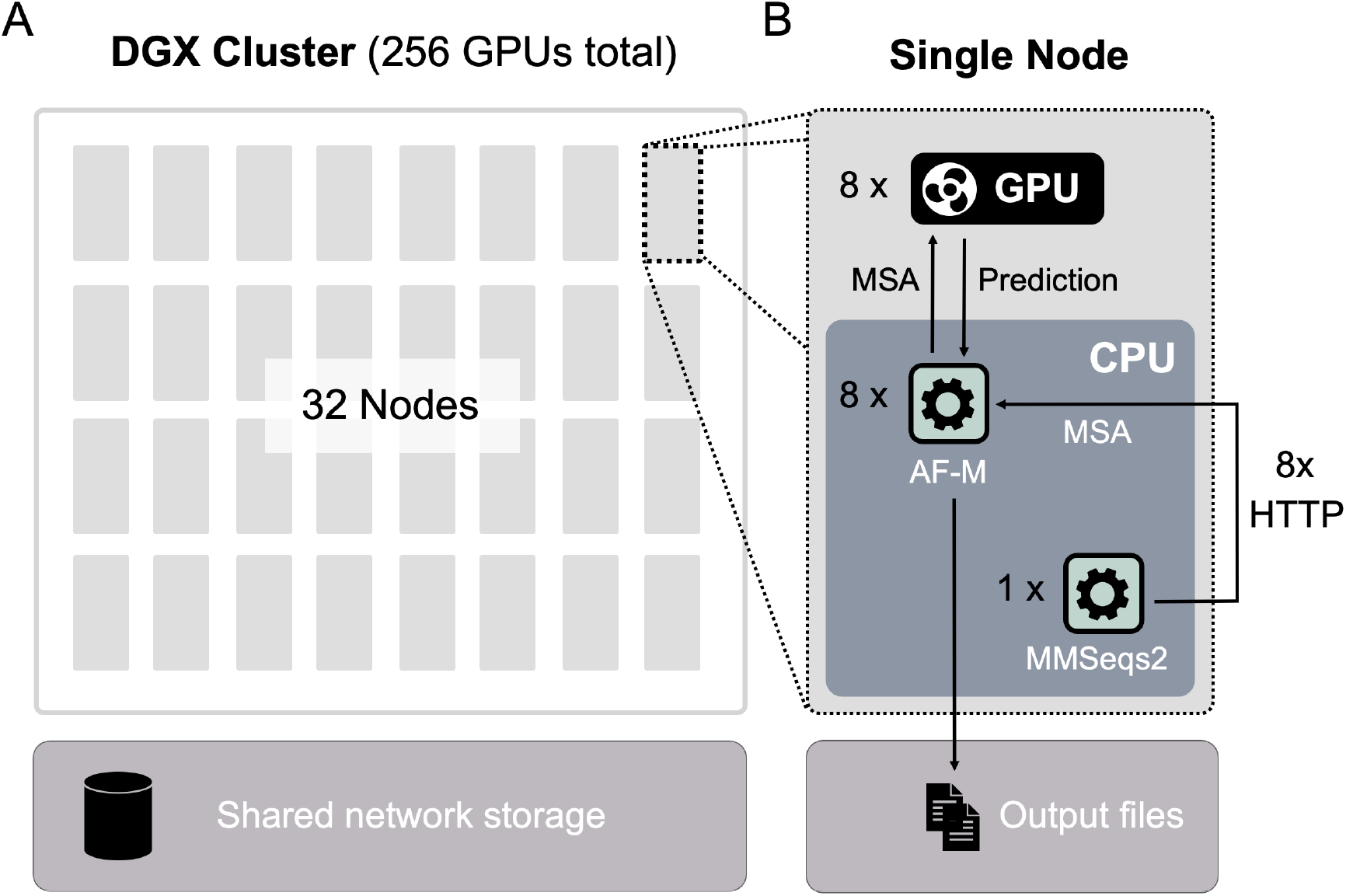
Computational architecture used for *in-silico* interaction screening. **A**. Schematic illustrating the architecture of the NVIDIA DGX cluster consisting of 32 nodes with 8 GPUs each, which was used to generate AF-M models for the 1.6 million KIRC-nominated protein pairs. **B**. A diagram displaying how various programs communicated within each node of the cluster, which operated independently and was supplied with a series of protein pairs to model. These pairs were split across the node’s 8 GPUs, and multiple sequence alignments (MSAs) were supplied via HTTP requests to a node-specific instance of the MMSeqs2 server software running on the node’s 8 CPUs. Once generated, predictions were deposited to a shared file system.

**Figure S2:**
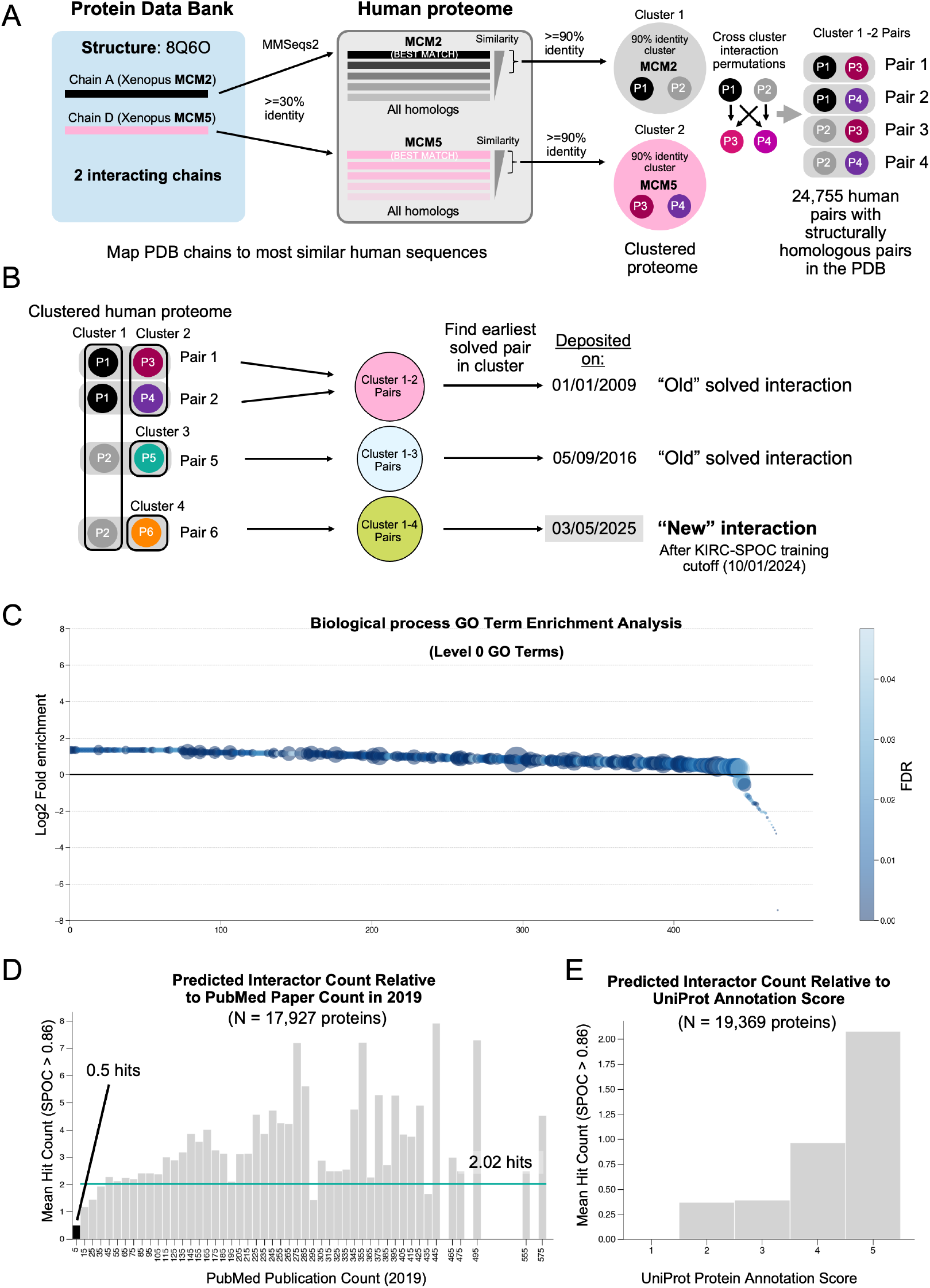
Analyzing the classes of proteins identified via *in-silico* screening. Schematic illustrating how protein interactions in the Protein Data Bank (PDB) were mapped to homologous human pairs. Schematic demonstrating how interactions were clustered by sequence and subsequently assigned a first PDB deposition date. **C**. A scatter plot of all enriched (>1) or de-enriched (<1) bioprocess GO terms (Level 0/Top level terms) associated with the 8,199 proteins in the high-confidence 16k set. The y axis is the log2 of the fold enrichment, and all terms are ranked high to low, left to right. Circle size is proportional to the number of proteins associated with that term, and circle fill color is based on the FDR associated with term enrichment analysis. **D**. Histogram of the mean number of high-confidence SPOC hits (>0.86) versus the PubMed publication count for 17,927 proteins across the screen. The horizontal teal line is the mean number of hits for all proteins with 10 or more publications in PubMed^21^. **E**. Histogram of mean number of high-confidence SPOC hits (>0.86) versus the UniProt Protein Annotation score for 19,369 proteins across the screen.

**Figure S3:**
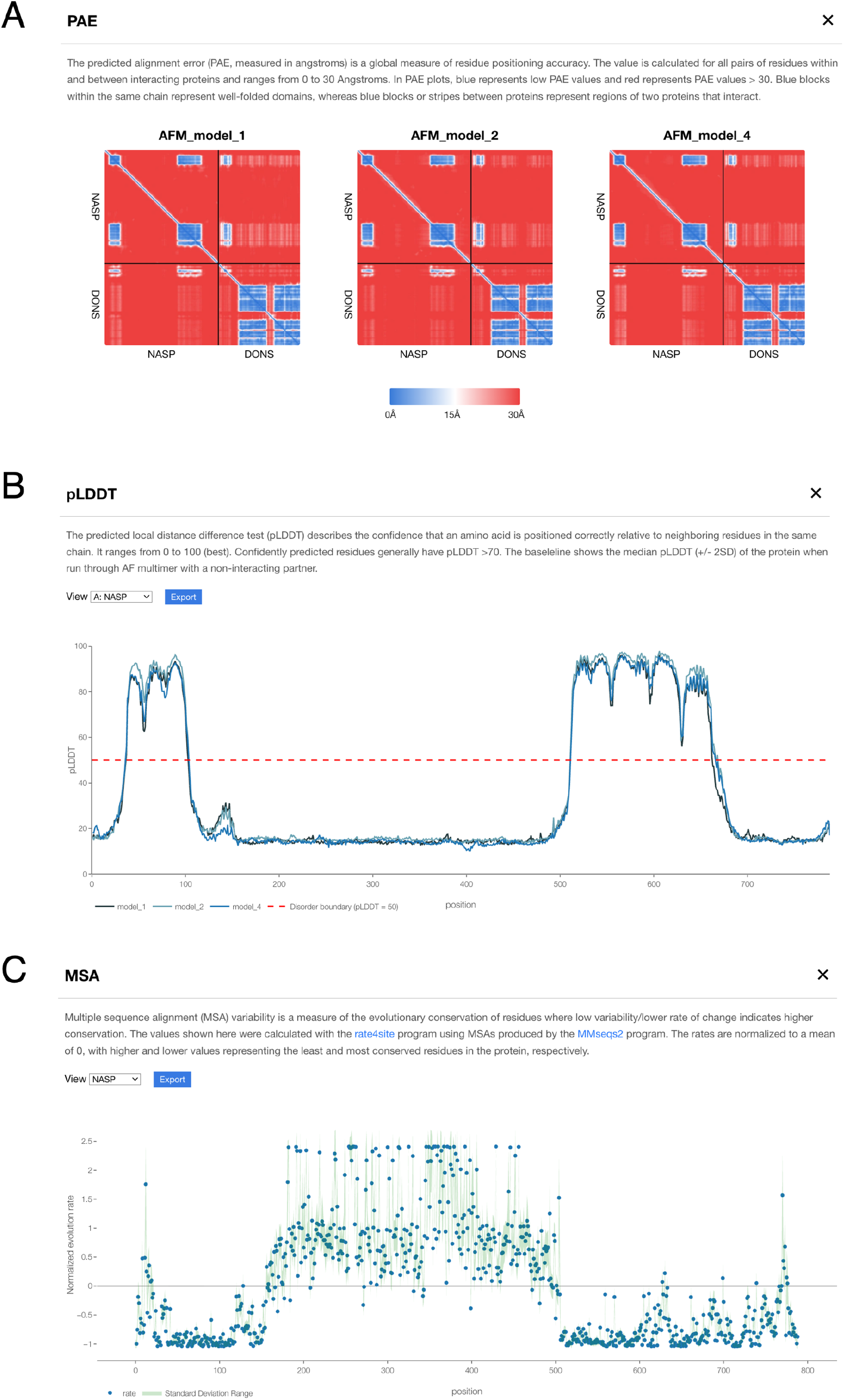
Additional information available at predictomes.org. Screenshot from a predicted interactor’s information page at predictomes.org showing interactive **(A)** Predicted Aligned Error (PAE), **(B)** Predicted Local Distance Difference Test (pLDDT), and **(C)** Multiple Sequence Alignment (MSA) plots.

**Figure S4:**
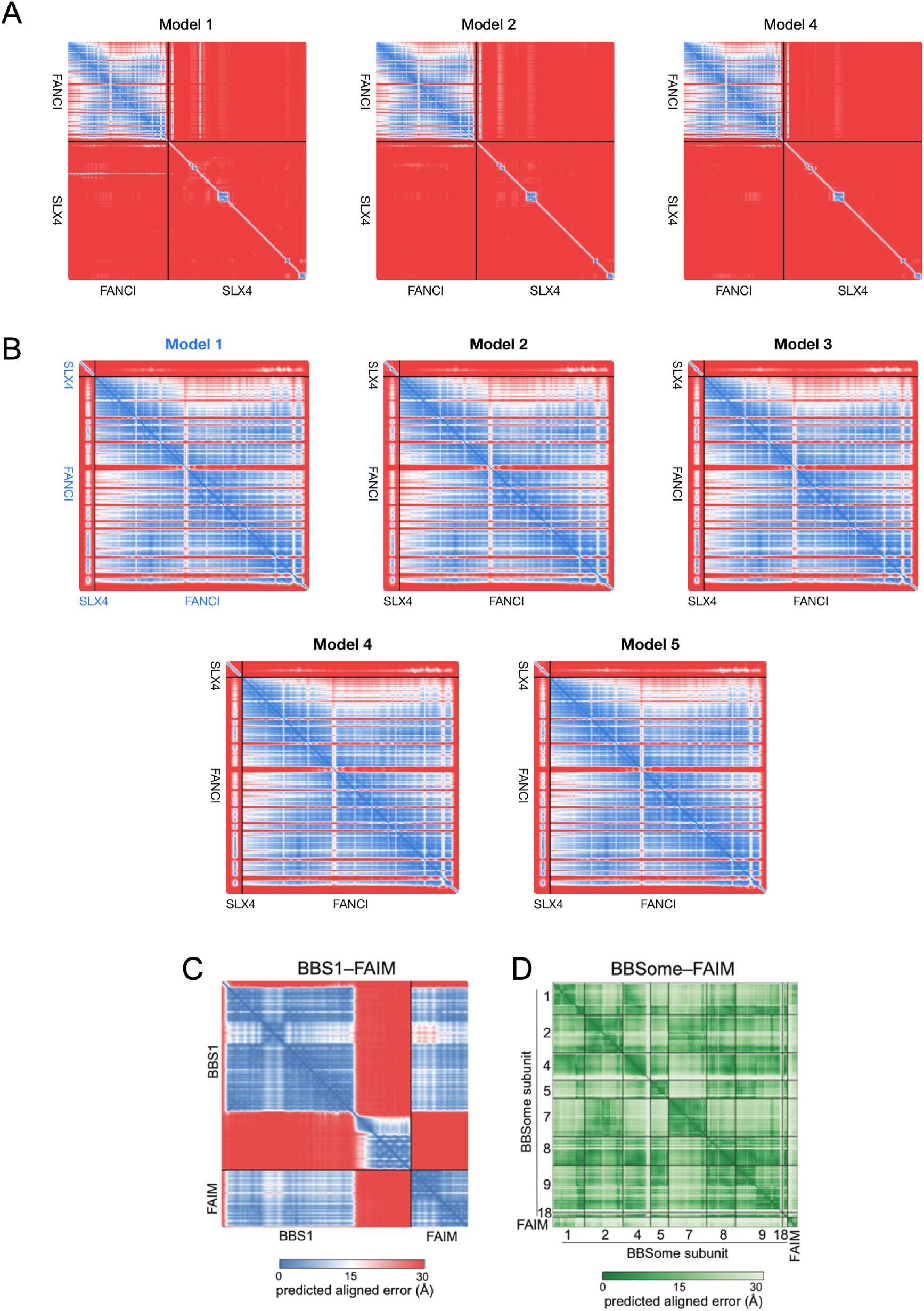
Predicted aligned error (PAE) plots supporting novel hypotheses. **A**. PAE plots depicting the FANCI–SLX4 interaction for all three predicted models retrieved from predictomes.org. The color scale indicates the per-residue predicted error, with blue representing low error and red representing high error. **B**. PAE plots showing five predicted models of the interaction between FANCI and SLX4 residues 1-100. **C**. PAE plot of a single model for the BBS1–FAIM interaction retrieved from predictomes.org. The PAE plots for the other four models look similarly confident. **D**. PAE plot of the BBSome–FAIM complex generated by AlphaFold3. In the color scale, darker green represents lower per-residue predicted error.

**Figure S5.**
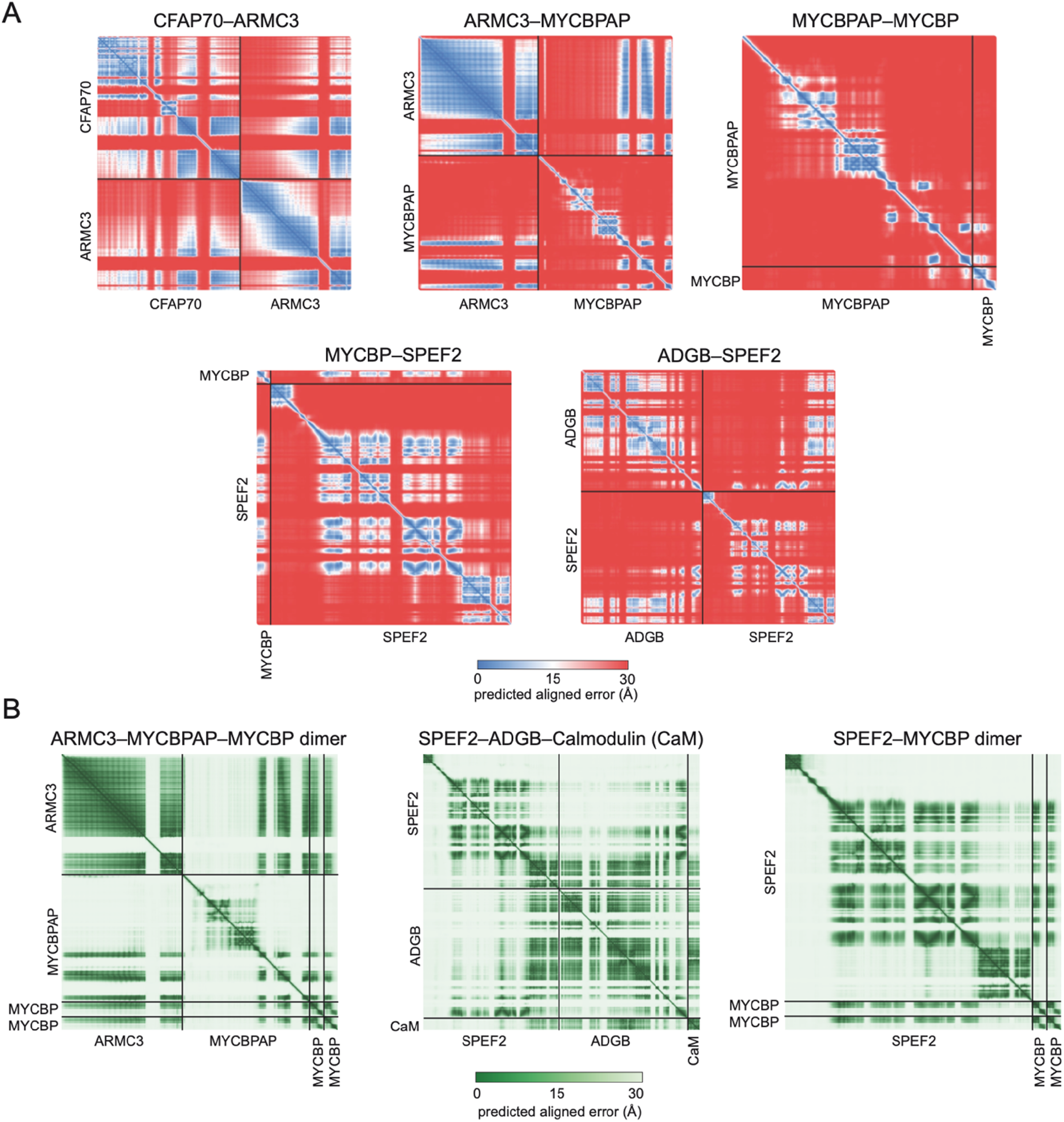
Predicted aligned error (PAE) plots for interactions involving central apparatus proteins. **A**. PAE plots for pairwise predictions using human protein sequences retrieved from predictomes.org. Only the PAE plot for model 1 is shown for each prediction, but the other PAE plots look similarly confident. The color scale indicates the per-residue predicted error, with blue representing low error and red representing high error. **B**. PAE plots for multi-subunit predictions using AlphaFold3 with mouse protein sequences. The color scale represents the expected position error, with darker green indicating higher confidence.

**Figure S6.**
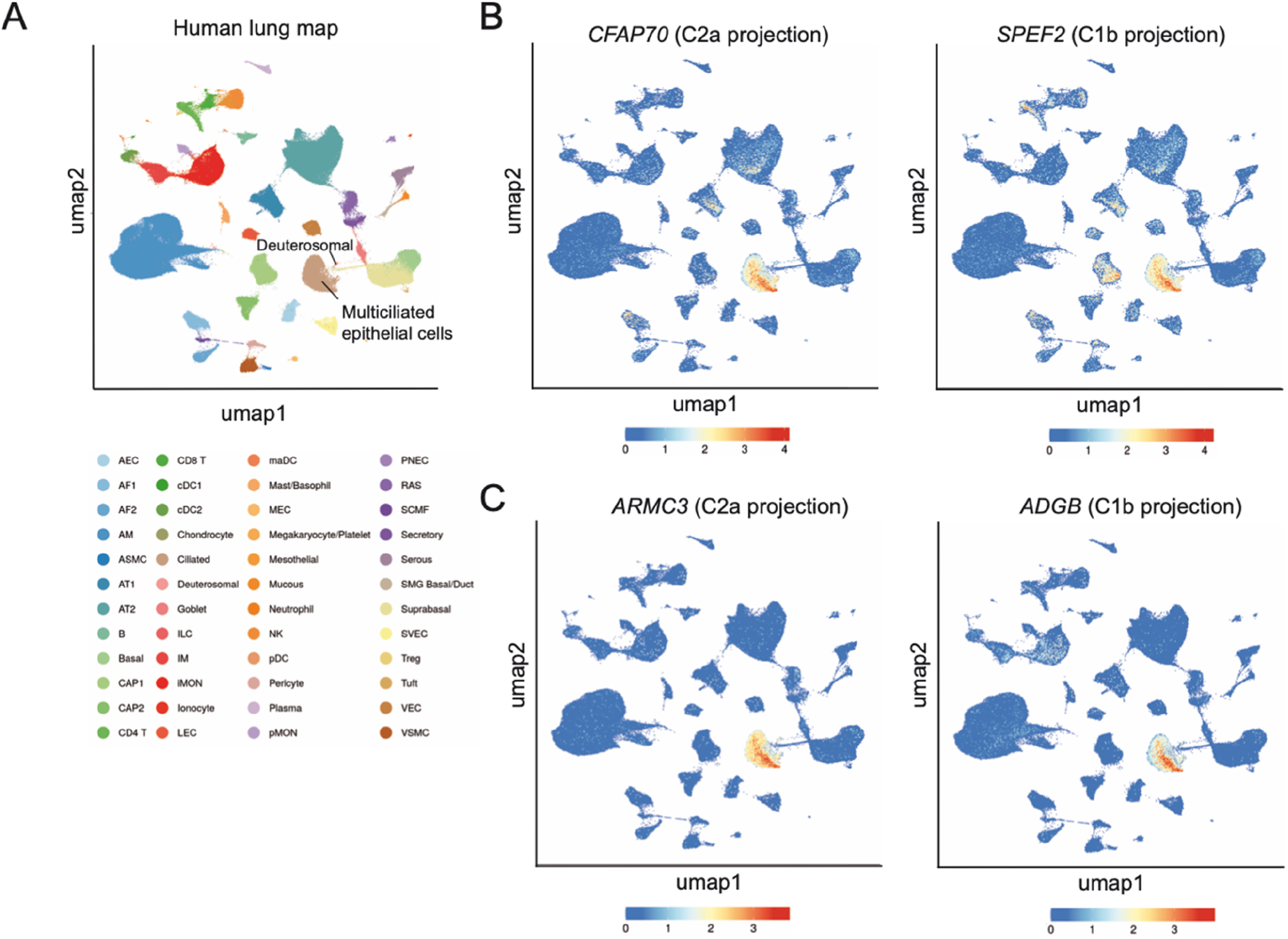
*ARMC3* and *ADGB* are expressed in the multiciliated cells of the lung. **A**. Uniform Manifold Approximation and Projection (UMAP) plot showing clustering of major human lung cell types generated by integrating 148 single-cell or single-nucleus RNA-seq datasets from 104 human lung donors^49^. Cells are colored according to their predicted identities. Labeled are multiciliated epithelial cells and their precursor deuterosomal cells. **B**. UMAP feature plots displaying expression of *CFAP70* and *SPEF2*, with color intensity indicating normalized gene expression (blue: low, red: high). Expression is mostly restricted to the multiciliated epithelial cells, consistent with their gene products being components of the central apparatus of motile cilia. **C**. UMAP feature plots displaying the expression of *ARMC3* and *ADGB*. Both genes have expression profiles that closely resemble *CFAP70* and *SPEF2* as shown in panel B.

**Figure S7:**
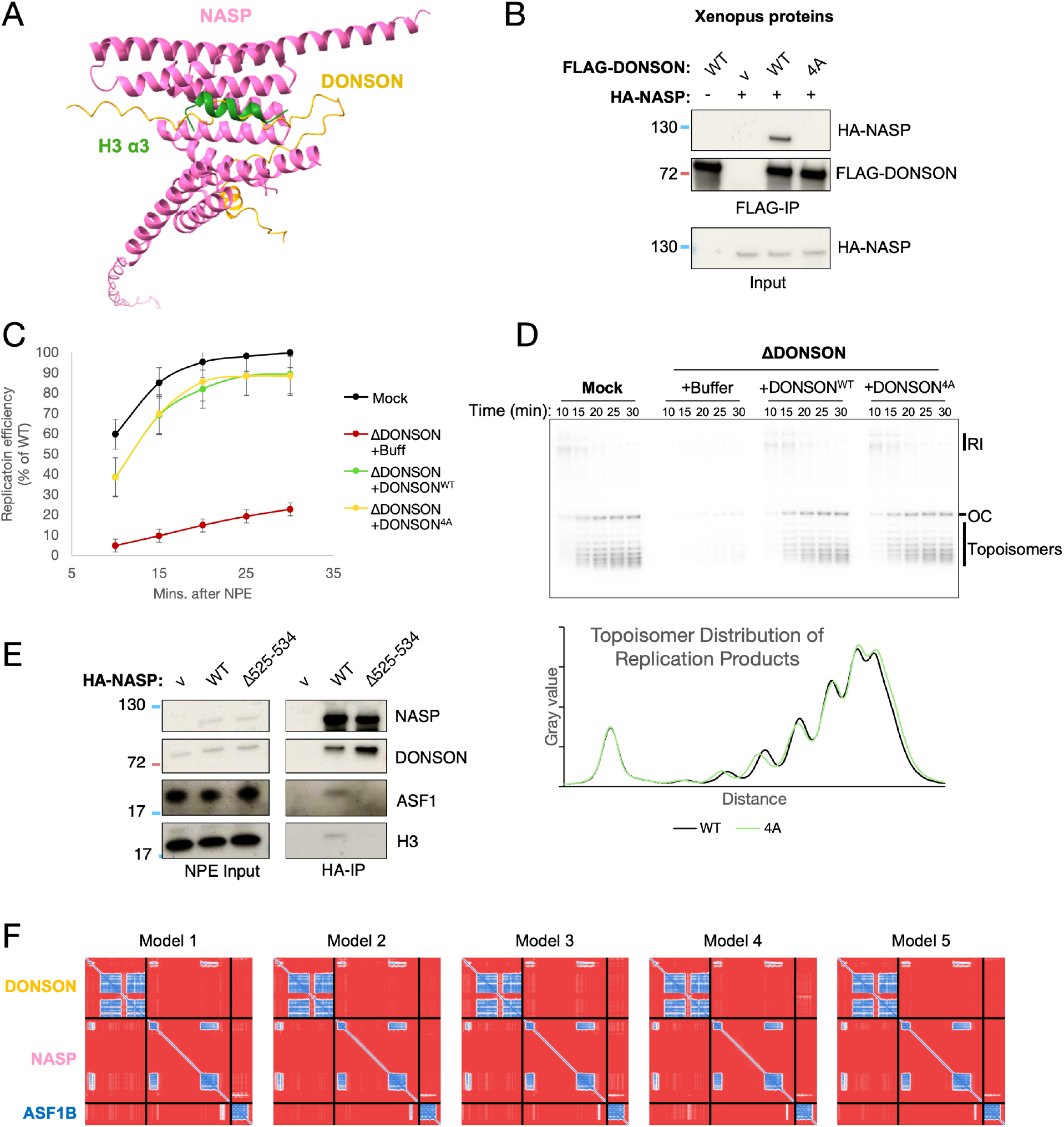
Mechanistic insights into DONSON’s interaction with NASP. **A**. Superimposition of the AF-M predicted DONSON–NASP complex with the NASP–H3 crystal structure (PDB: 7V1L). The overlay illustrates that the DONSON-binding surface on NASP overlaps with the H3-H4-binding interface, indicating that these interactions are mutually exclusive. **B**. Co-immunoprecipitation of *Xenopus* NASP with the indicated FLAG-DONSON variants. Same experiment as in Figure 6D, except using *Xenopus* NASP. **C**. The interaction of DONSON with NASP is not required for efficient DNA replication. DONSON was immunodepleted from egg extracts (NPE), and recombinant CDK2–Cyclin E1 (to restore efficient replication due to partial CDK2–Cyclin E co-depletion) and buffer or the indicated mutants were added back. Licensing and replication initiation were carried out in the presence of radioactive dATP, and replication efficiency was measured by running replication products on native agarose gels and performing autoradiography (see Methods). Data represent mean ± SD (n = 3). **D**. Same experiment as in (C), but plasmids were separated on native gels containing 1 µM chloroquine to resolve plasmid topoisomers, and the autoradiograph of the gel is shown (top), together with a density trace through each lane (bottom), to indicate that there is no difference in the degree of plasmid supercoiling in the different conditions where replication was observed, consistent with there being no major defect in replication-coupled chromatin assembly. **E**. Independent replicate of the data shown in Figure 6F. **F**. Predicted aligned error (PAE) plots of the AF-M predicted DONSON–NASP–ASF1B complex. The color scale indicates the per-residue predicted error, with blue representing low error and red representing high error.

**Figure S8:**
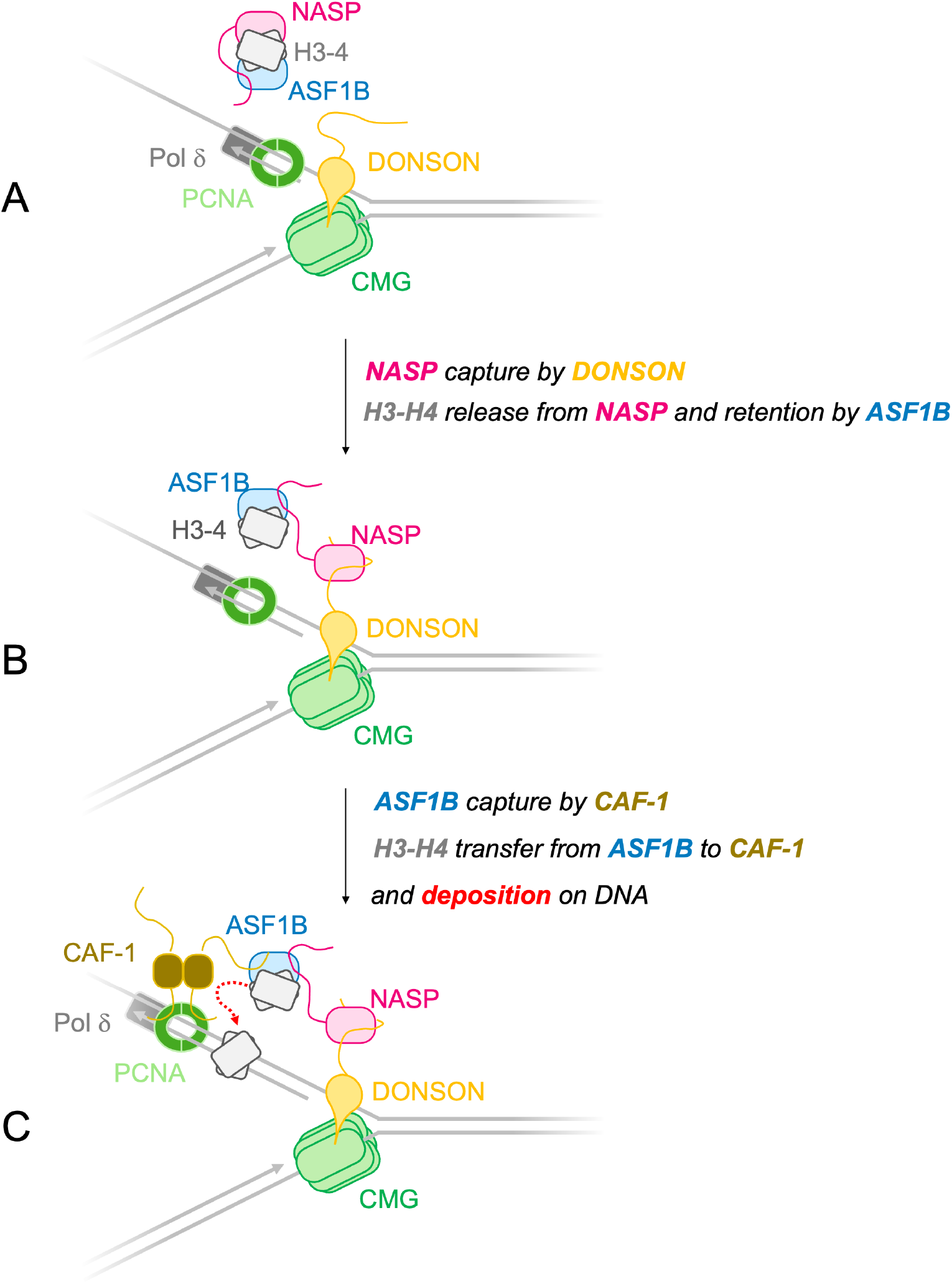
Hypothetical model of DONSON’s role in chromatin assembly. Given the new protein-protein interactions reported here (DONSON–NASP and NASP–ASF1B) and the similar effects of DONSON, NASP, and CAF-1 depletion on fork progression, we hypothesize that these proteins cooperate in replication-coupled chromatin assembly, according to the following mechanism. **A**. Our data indicate that NASP and ASF1B bind cooperatively to H3-H4 because a NASP mutant that disrupts the newly identified NASP–ASF1B interaction also no longer binds histone H3. We further propose that DONSON interacts with the replisome through previously described interactions with GINS and MCM3. **B**. We propose that DONSON’s N-terminal region captures NASP. Because DONSON and histones bind the same surface on NASP, DONSON binding triggers histone transfer from NASP to ASF1B. Consistent with this model, NASP mutants that lose the interaction with histones exhibit a modest but reproducible increase in binding to DONSON (**Figures 6F and S7E**). The new interaction detected between NASP and ASF1B indicates that the ASF1B–H3-H4 complex might remain associated with NASP after DONSON binding. **C**. The known interaction between CAF-1 and ASF1B facilitates histones transfer from ASF1B to CAF-1, followed by deposition on DNA. We speculate that replication-coupled chromatin assembly remains intact in egg extracts when the DONSON–NASP interaction is disrupted because the large pool of free histones in this system bypasses the need for this pathway. Alternatively, direct interaction with CAF-1 or DONSON–NASP might represent redundant mechanisms for ASF1B–H3-H4 complexes to be concentrated near CAF-1.

